# Dynamic and task-dependent decoding of the human attentional spotlight from MEG

**DOI:** 10.1101/2025.10.23.684150

**Authors:** Maryam Mostafalu, Tommy Clausner, Maxime Ferez, Danila Shelepenkov, Sébastien Daligault, Denis Schwartz, Jérémie Mattout, Suliann Ben Hamed, Mathilde Bonnefond

## Abstract

Attention is a fundamental mechanism enabling the brain to overcome its limited capacity for parallel processing. In non-human primates, invasive electrophysiology has shown that attentional selection operates rhythmically, primarily within the alpha (∼8–12 Hz) and theta (∼4–5 Hz) bands. Whether such finely resolved control signals can be captured non-invasively in humans, and how they adapt to changing task demands, remains unclear. Using high-precision magnetoencephalography (MEG) combined with machine learning, we decoded the spatial locus of covert attention in humans performing three variants of a spatial cueing task that manipulated cue validity as well invalid trial switching rules.

Spatial attention could be decoded from whole-brain MEG activity at both static and time-resolved scales, with accuracies significantly above chance (N = 30). Decoding performance decreased as cue validity was reduced, indicating that task structure shapes attentional engagement. Analysis of decoding trajectories revealed rhythmic fluctuations at ∼8–12 Hz across all tasks, demonstrating alpha-band sampling of attention. Pre-target attention became increasingly focused on the cued side, especially in the 100% Valid condition, consistent with proactive orienting. Furthermore, individual and task-specific differences in decoding strength correlated with task-variations in behavioral performance, linking the accuracy of neural attention codes to both discrimination accuracy and reaction time.

These findings demonstrate that MEG can non-invasively capture dynamic, task-dependent fluctuations in spatial attention that parallel those observed in non-human primates. They reveal that attentional demands reshape the neural code for attention, modulate rhythmic sampling, and influence behavioral efficiency. This work bridges invasive primate and non-invasive human research and establishes MEG-based decoding of attention as a promising tool for mechanistic and clinical applications, including neurofeedback and attention-related interventions.

## Introduction

The brain’s processing bandwidth is inherently limited, necessitating mechanisms that prioritize information. Selective attention fulfills this role by biasing neural resources toward behaviorally relevant inputs, thereby supporting adaptive perception and decision-making. Accordingly, attention has become a major focus of research in both humans and animal models, and its neural implementation remains a central target for basic and clinical neuroscience.

Classic theories of attention describe selection as stable in space and time. In contrast, recent behavioral and neural results show that attention samples the environment rhythmically. The frequencies most often reported are in the alpha frequency band, ∼8–12 Hz, and the theta frequency band, ∼4–5 Hz (Crouzet and VanRullen, 2017; Fiebelkorn et al., 2013; Gaillard et al., 2020; Kienitz et al., 2018; Landau and Fries, 2012; Spyropoulos et al., 2018; VanRullen, 2013, 2016) The specific frequency observed in a given setting depend on the task difficulty and spatial organization of the task, supporting the idea that rhythmic attention subserves task-relevant information prioritization (Holcombe and Chen, 2013; Michel et al., 2020; VanRullen, 2016). Several models have been proposed within the rhythmic theory of attention (Gaillard and Ben Hamed, 2022; Hamed, 2025) . One model suggests that attention blinks from one task-relevant spatial location to another. VanRullen and colleagues (2018) proposed that perception proceeds in discrete cycles rather than continuously, with attention operating as a temporal filter that rhythmically modulates perceptual gain. These oscillatory cycles, typically in the theta or alpha range, are proposed to organize alternating phases of sensory enhancement and suppression, thereby coordinating exploration of new information and exploitation of task-relevant inputs (Fiebelkorn and Kastner, 2019; VanRullen, 2016, 2018) and changing the resolution of ongoing information processing (Michel et al., 2020). An alternative model proposes that attention does not sample space through discrete perceptual cycles but rather explores it continuously through rhythmic attentional saccades (Gaillard et al., 2020). Using real-time decoding of prefrontal activity in macaques, they showed that the attentional spotlight follows continuous trajectory across visual space, with sharp directional transition, a.k.a. attentional saccades, at ∼8 Hz within the alpha band. Even though the task designated a single behaviorally relevant location, the decoded spotlight was also found at other spatial positions, eliciting inappropriate responses to task-irrelevant stimuli (Di Bello et al., 2022; Gaillard et al., 2020). This rhythmic processing of space is proposed to alternate between exploitation of the task-relevant location and exploration of surrounding space, providing a dynamic balance between focused goal-directed and distributed stimulus-driven processing (Di Bello et al., 2022; Gaillard et al., 2020), possibly to allow rapid responses to unexpected events.

Previous work (Astrand et al., 2016; De Sousa et al., 2021) developed a precise method that enables a spatially and temporally resolved decoding of the locus of attentional selection, i.e. the attentional spotlight. This methodological advance has deepened our understanding of the neural bases of attention (Amengual et al., 2022; Astrand et al., 2020; Di Bello et al., 2022) and the rhythmic nature of attention (Gaillard et al., 2020). The method reads out the position of the attentional spotlight along the x and y axes with high spatial and high temporal precision, using signals from the prefrontal cortex of macaques, and relying on either multi-unit activity or local field potentials (LFP). This high resolution decoding has proven instrumental in revealing the rhythmic properties (Gaillard et al., 2020) and state-dependent dynamics (Amengual et al., 2022) of attention, as well as in uncovering how prefrontal networks flexibly implement both proactive and reactive mechanisms of visual suppression (Di Bello et al., 2022), and how these mechanisms dynamically interact with task-identity (Mouille et al., 2025).

A key open question is how top-down control flexibly governs the dynamic spatial orientation of attention. Prior studies have typically used cues with 100% validity, ensuring full predictability of target location and thus promoting proactive orienting toward a single spatial goal. Under such conditions, attention can remain stably focused, and rhythmic fluctuations primarily reflect internal cycles of exploration and exploitation within the cued hemifield. However, real-world environments rarely afford such stability. When predictive information becomes uncertain, efficient behavior requires a more dynamic balance between maintaining focus on the most probable location and monitoring alternative, potentially relevant positions. To date, the precise neural mechanisms supporting this task-related flexibility remain unclear. Answering these questions in macaques poses practical challenges. Invasive electrophysiology is technically demanding and logistically intensive: animals must be trained on several task sets, a process that is both time-consuming and resource-heavy and may introduced uncontrolled for experimental biases. Moreover, resolving how task structure shapes attention would ideally require recordings from multiple brain areas, yet implant-based approaches are necessarily restricted to a small set of regions of interest. Taken together, these constraints slow progress toward a fuller account of attentional mechanisms and complicate the translation of findings to human applications.

In the present study, we combine high-precision magnetoencephalography (MEG) with decoding approaches inspired by macaque work to examine whether the rhythmic attentional spotlight can be reconstructed from human brain activity and how its spatial distribution, frequency and correlation with behavior are shaped by task structure. Unlike other whole-brain approaches in humans (e.g., EEG or fMRI), MEG primarily captures postsynaptic dendritic currents, signals closely related to the local action and field potentials used for decoding in nonhuman primates, while offering high temporal resolution together with meaningful spatial resolution. This correspondence supports the translation of decoding strategies from macaques to humans. Using this framework, we decoded spatial attention across two MEG sessions and three different spatial cueing tasks with variable cue validity rules. The results that spatial attention can be decoded from whole-brain MEG activity with high temporal precision, mirroring the fine-grained intracortical findings previously obtained in macaques. Manipulations of task structure significantly reshaped both decoding performance and the spatial distribution of pre-target attention, demonstrating that human cognitive flexibility modulates the underlying attentional code. Moreover, decoding strength correlated with behavioral performance (accuracy and reaction time) in a task-specific manner, indicating that the neural representation of attention dynamically adapts to the cognitive and predictive demands imposed by each task. Finally, the decoded attentional signal oscillated at ∼8–12 Hz across all task variants, confirming that rhythmic sampling constitutes a core organizing principle of human attention, with task-dependent variations in oscillatory strength.

## Methods

### Participants

Forty-five healthy, right-handed volunteers (19 males, 26 females, aged 26.68±0.61 years) with normal or corrected-to-normal vision and no history of neurological or psychiatric disorders were included in the study. Five participants withdrew from the study prior to the MEG sessions reported here due to COVID-19 related circumstances. Additionally, five participants were excluded because of poor behavioral performance, and five others were excluded due to excessive technical problems with eye-tracking data recording. This resulted in a final sample of 30 participants (12 males, 18 females, aged 27.1±0.72 years). All participants provided written informed consent approved by a French Research Ethics Committee (reference number NCT04175119) in accordance with the Declaration of Helsinki.

### General Procedure

Participants completed six sessions (MRI/fMRI, EEG, and four MEG sessions) over a maximum three-month period. At each session, informed consent and a COVID-19 related questionnaire were collected, along with additional questionnaires regarding recent sleeping behavior and lifestyle. This paper focuses on the last two MEG sessions examining covert spatial attention.

### Neuroimaging Acquisitions

#### MRI Data Acquisition

We acquired structural MRI data using a 3T Magnetom Prisma scanner (Siemens Healthcare, Erlangen, Germany) with the body coil for radio-frequency (RF) transmission and a standard 32-channel RF head coil for reception. Two scans were performed; however, only the one relevant to the present study is described below. **T1-weighted scan**: A 3D spoiled fast low angle shot (FLASH) sequence with 1 mm isotropic resolution, field-of-view of 256×256×192 mm along the anterior-posterior, head-foot, and right-left directions, respectively. The repetition time was 7.96 ms with an excitation flip angle of 12°. A high readout bandwidth (425 Hz/pixel) preserved brain morphology without geometric distortions. This scan, which included the participant’s neck, was used for individual head-cast creation. Acquisition time was 3 min 42 s using a 32-channel head coil without padding or headphones to avoid scalp distortions.

#### MEG Recording

Before MEG measurement, participants received instructions and completed a practice block outside the scanner. Once comfortable with the task, they were fitted with an individually-designed head-cast allowing to reduce head-movements within the MEG helmet, three fiducial marker coils were embedded into the surface of the head-casts at nasion, left and right ear canals to monitor the subject’s relative head motion, and they entered the scanner room. MEG data were recorded in a standard magnetically shielded room consisting of two layers of μ-metal and one layer of copper (Vacuumschmelze, Hanau, Germany), without active shielding. The residual background magnetic field averaged approximately 20 nT. We used a 275-channel SQUID-based axial gradiometer system (CTF MEG Neuro Innovations Inc., Port Coquitlam, BC, Canada) at 1200 Hz sampling rate. Eye movements were simultaneously monitored using an EyeLink eye tracker system (SR Research Ltd.) at a 1000 Hz sampling rate to ensure fixation stability.

### Experimental Design

Participants performed three different versions of a covert spatial attention task across two MEG sessions. Several blocks of trials were collected for each task version. We will first describe trial structure, then task versions, then block organization (**Figure 1A-E)**.

**Figure 1.**
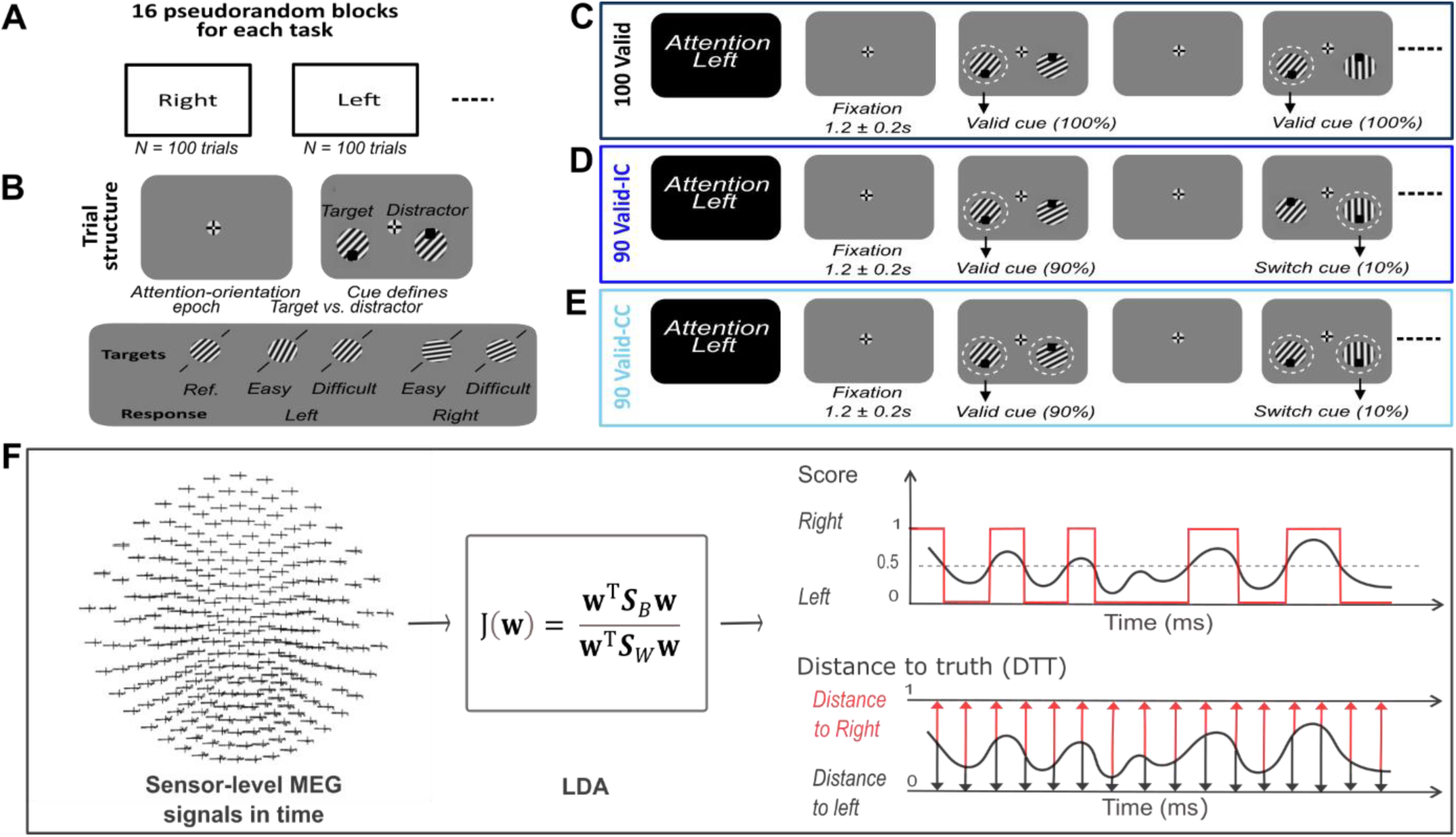
Task design and experimental conditions. **(A) Block structure.** Participants performed covert spatial attention tasks across 16 pseudorandomized blocks per condition. Each block consisted of 100 trials in which participants were instructed to attend either the left or right visual field. The cued side remained constant within a block but was counterbalanced across blocks (50% left, 50% right). **(B) Trial structure.** Within a block, each trial began with a central fixation cross (duration: 1.2 ± 0.2 s), followed by simultaneous presentation of two oriented gratings at 3° eccentricity in the left and right bottom visual fields. Gratings varied in orientation relative to an imaginary diagonal (45° from vertical). Each trial was categorized as Easy or Difficult, depending on the angular deviation of the grating lines relative to the diagonal (larger deviations for Easy trials, smaller deviations for Difficult trials). Participants had to identify whether the target grating deviated clockwise or counterclockwise relative to this diagonal. Each grating included a small black square either at the bottom or top indicating target and distractor respectively (see Panels C–E). **(C) 100 Valid cue Task.** In this task, the cue indicating the attended side was always valid. The position of the black squares remained unchanged across all trials within a block (bottom on the attended grating, top on the unattended grating). Participants could rely entirely on the initial instruction to maintain attention on the cued side. **(D) 90 Valid-IC Task.** In this task, cues were valid in 90% of trials. On 10% of trials (invalid), the black square on the attended grating flipped vertically from bottom to top, while the square on the unattended grating flipped from top to bottom, signaling that participants should shift attention to the opposite side. This manipulation required rapid attentional reorienting in response to an explicit switch cue. **(E) 90 Valid-CC Task.** In this task, cues were also valid in 90% of trials. On invalid trials (10%), only the black square on the unattended grating flipped from top to bottom, requiring participants to monitor the unattended side throughout the block and detect these occasional changes. This manipulation required distributed monitoring of both attended and unattended locations. **(F) Overview of decoding analysis pipeline and calculation of Distance to Truth (DTT). Left panel.** MEG signals were extracted from visual channels in the interval from –500 ms to +300 ms relative to stimulus onset for all trials and served as input features to the decoding model. Each trial was labeled according to the attended side (0 = left, 1 = right), forming the ground truth labels used for classification. **Middle panel. Linear Discriminant Analysis (LDA) decoding.** At each time point, a Linear Discriminant Analysis (LDA) classifier was trained to discriminate left-attended versus right-attended trials. LDA generated a continuous output score ranging from 0 to 1, indicating the classifier’s confidence in the right-attended label. **Right panel, top. Decoding accuracy.** Decoding accuracy (red) was computed as the proportion of trials for which the predicted label (thresholded LDA output, black curve) matched the true label, relative to the total number of trials. This yielded time-resolved accuracy traces quantifying classifier performance across the epoch. **Right panel, bottom. Distance to Truth (DTT) metric.** For each trial and time point, the classifier’s continuous output score was compared to the true label, and the absolute difference was calculated as the Distance to Truth (DTT). Lower DTT values indicate classifier outputs closer to the actual cued location, reflecting more precise attentional selection.

#### Trial structure

Stimuli were projected to a screen of ≈ 50 cm distance to the subject’s eyes. Throughout the experiment, participants were instructed to maintain central fixation, shifting only their attention (not their gaze). A central fixation target designed to maximize fixation performance (Thaler et al., 2013) was positioned centrally. Participants were instructed to blink only after responding to avoid blinking before and during stimulus presentation. On each trial (**Figure 1B**), after a fixation epoch of 1.2 ± 0.2s, two gratings appeared simultaneously at 3° eccentricity on the left and bottom visual field (3 degrees away from center in the x and y axis). Participants judged whether the orientation of the grating lines on the cued side deviated clockwise or counterclockwise from an imaginary diagonal, responding with their right index or middle finger. We manipulated the predictability of grating orientation using a transition matrix. For the attended side only, a given grating would predict one specific grating with 80% likelihood, another grating with the same orientation category with 10% likelihood, and the two gratings with opposite orientation with 5% each (results not presented here).

#### Task versions

Task versions were defined as follows. Each grating contained a small black square (either on top or bottom) that served as a potential attention-switching cue. On the valid trials of each block, squares on attended gratings were positioned at the bottom while squares on unattended gratings were at the top. We implemented three experimental conditions or tasks that manipulated cue validity. How these squares were positioned on the next trials defined the task version subjects were presented with as follows. **1) 100 Valid**: 100% valid cues, where targets always appeared on the initially cued side (**Figure 1C**). The squares remained in their initial positions throughout the block. **2) 90 Valid-IC**: 90% valid cues, with invalid trials (10%) signaled by the black square flipping vertically (from bottom to top) on the attended stimulus, requiring participants to shift attention to the opposite side (**Figure 1D**). Simultaneously, the square on the unattended side flipped from top to bottom. **3) 90 Valid-CC**: 90% valid cues, with invalid trials (10%) signaled by the black square flipping vertically (from top to bottom) on the unattended side, requiring participants to monitor the unattended side throughout the block for possible switches (**Figure 1E**). The *90 Valid-IC* and *90 Valid-CC* conditions were presented in different sessions (counterbalanced across participants) to avoid confusion.

#### Block organization

Each block began with a 5-second baseline period where participants fixated centrally and avoided blinking. They then received explicit instructions about the attention condition (attend left or right) and cue validity (always or mostly valid, **Figure 1A**). Each trial began with a central fixation point, followed by simultaneous presentation of both gratings after a variable interval (1.2 ± 0.2s). Participants responded, using a two-button button box, as quickly and accurately as possible using their right-hand index finger for counter-clockwise deviations and middle finger for clockwise deviations. After response, a new trial began. Each block comprised 100 trials, after which participants received performance feedback. Feedback on eye fixation quality throughout the block was additionally provided.

The target attention side remained constant within blocks but was counterbalanced (50:50) between blocks. In the first session, participants completed 32 blocks of 100 trials each: 16 blocks of 100 VALID condition and 16 blocks of either *90 Valid-IC* or *90 Valid-CC* (counterbalanced across participants). Importantly, the first 4 blocks were always *100 Valid* condition to facilitate implicit learning of the transition matrix. In the second session, participants completed 16 blocks of the alternate V90 condition (either *90 Valid-IC* or *90 Valid-CC*, depending on what they completed in the first session). In total, participants performed 1600 trials per condition.

#### Stimuli

Stimuli consisted of four gratings with distinct orientations for each participant (**Figure 1B**). The gratings presented 12 cycles across and were 3.5° visual angle in diameter. The grating lines were oriented relative to an imaginary diagonal that could be either left or right with a 45° angle relative to the vertical axis. The reference diagonal remained consistent for each participant throughout all experimental sessions but was counterbalanced across participants. Deviations from the diagonal could be achieved by clockwise or counterclockwise rotation of the grating. The diagonal itself was not visible.

During the first MEG session, 34 participants completed a staircase procedure in which the deviation from the diagonal was varied between ±1° and ±44°. Stimuli were presented unilaterally in the bottom right of the screen at 3° distance. The deviation angle that yielded 80% correct responses was selected as a reference for subsequent experiments. Angles derived from the "far" condition (generally larger) were used as "easy" trials (mean angle 18.35° ± 2.16), while those from the "close" condition (generally smaller) were used as "difficult" trials (mean angle 12.42° ± 1.65). For 11 (6 of which are included in this work) participants, we could not perform the staircase procedure for technical reasons, we therefore used default angles of 30° and 15° for easy and difficult conditions, respectively.

### Data preprocessing

Data were preprocessed using the FieldTrip toolbox in MATLAB. Trials were extracted from raw MEG recordings, and automatic artifact rejection was applied to eliminate prominent signal artifacts. Eye-tracking data were analyzed to identify and remove trials containing blinks at any point or saccades exceeding 1.5° from central fixation. We specifically excluded trials containing blinks or saccades within the critical interval from 500 ms pre-stimulus to stimulus onset. Only trials with correct responses were included in further analyses, as correct responses indicated effective attentional allocation to the cued side.

### Behavioral analysis

Behavioral analyses were performed on reaction time (RT) and response accuracy data for each participant and task. Only correct-response trials were included in the RT analysis. Trials were categorized as *easy* or *difficult* based on the target grating’s orientation degree relative to the diagonal, and as *congruent* (CS) or *incongruent* (IS) according to whether the target and distractor orientations matched.

#### Valid trials

For valid trials, we used a three-way repeated-measures ANOVA with within-subject factors Task (*100 Valid, 90 Valid–IC, 90 Valid–CC*), Difficulty (Easy, Difficult), and Congruency (CS, IS). The dependent variable was the median RT for each condition and participant. Separate ANOVAs were conducted for RT and accuracy. When assumptions of sphericity were violated, Greenhouse–Geisser corrections were applied. Follow-up pairwise comparisons were conducted using two-sided Wilcoxon signed-rank tests to assess the effects of Difficulty (Easy vs. Difficult) and Congruency (CS vs. IS) within each task.

#### Invalid trials

Invalid (switch) trials from the *90 Valid–IC* and *90 Valid–CC* tasks were analyzed separately to assess behavioral effects during attentional reorienting. The same three-way repeated-measures ANOVA structure was applied, with within-subject factors Task (*90 Valid–IC*, *90 Valid–CC*), Difficulty (Easy, Difficult), and Congruency (CS, IS), using median RTs (correct trials only) and accuracy as dependent variables. Post hoc Wilcoxon signed-rank tests were performed to evaluate pairwise comparisons for Difficulty and Congruency within each task and, where relevant, to explore task × factor interactions.

### Pre-target alpha-band lateralization across tasks

As a sanity check, we examined whether the data exhibited the expected prestimulus alpha-band lateralization in the occipito-parietal channels. We computed time–frequency representations (TFRs) of the raw MEG signal separately for trials in which attention was cued to the left versus the right visual field and then plotted their contrast (left – right) in the interval of 500ms before target onset to target onset. **Figure S1**, panel A and B correspond to the TFR of this contrast for left and right hemisphere channels respectively across all three tasks separately. Panel C demonstrates the *topoplot* of the contrast TFR in the interval of -100ms to 0 across frequency range of 11 to 13 Hz. This contrast revealed the canonical pattern of hemispheric alpha modulation: alpha power increased over posterior sensors contralateral to the distractor and decreased contralateral to the attended (target) side. This pattern confirms the expected prestimulus alpha lateralization associated with spatial attention.

### Decoding Spatial Attention

To decode the attended location (left vs. right) from MEG data, we applied two complementary Linear Discriminant Analysis (LDA) methods. MEG signals were extracted from visual channels in the interval from –500 ms to +300 ms relative to stimulus onset for all trials. A 50-ms sliding window was applied with a 5-ms step size, resulting in time-resolved data segments that served as input features to the decoding model. These served as input features to the LDA (**Figure 1F**, left panel). Each trial was labeled according to the attended side (0 = left, 1 = right), forming the ground truth labels used for classification. LDA identifies neural patterns distinguishing predefined classes by maximizing the ratio of between-class variance to within-class variance, formalized by:

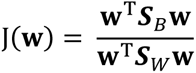

where **w** is the discriminant vector, ***S****_B_* denotes the between-class scatter matrix, and ***S****_W_* denotes the within-class scatter matrix (**Figure 1F**, middle panel).

#### Static Decoding

We computed mean neural activity for each MEG visual channel over four pre-stimulus windows (50 ms, 100 ms, 200 ms, and 300 ms before stimulus onset). For each window, we applied a Hann window by weighting the samples within the segment, normalizing the weights to sum to 1, and computing the resulting weighted mean amplitude per channel. For each participant and each time window, a linear discriminant analysis (LDA) classifier was trained to decode the attended side using these mean amplitudes as features. Decoding performance was calculated separately for each window length and each task condition (*100 Valid, 90 Valid-IC,* and *90 Valid-CC*).

#### Time-Resolved decoding

To examine the temporal dynamics of attentional allocation, we implemented a sliding window analysis using 50 ms windows shifted in 5 ms increments from –500 ms to +300 ms relative to stimulus onset. Within each window, a Hann window was applied to weight the samples before averaging, producing a weighted mean amplitude per channel per time point. These measures were used as features in an LDA classifier to generate continuous decoding accuracy time courses for each condition (*100 Valid*, *90 Valid-IC*, and *90 Valid-CC*). Decoding accuracy was computed as the proportion of trials for which the predicted label (thresholded score) matched the true label, divided by the total number of trials. This yielded time-resolved accuracy traces quantifying classifier performance across the epoch (**Figure 1F**, top right panel). Statistical significance at each time point was determined by comparing observed decoding accuracies against the null distribution from permutation tests, with multiple comparison correction applied using Bonferroni adjustment.

### Distance to Truth (DTT) and Spatial Distribution of Attention

We quantified moment-by-moment attentional alignment using the Distance to Truth (DTT), defined as the absolute difference between the classifier’s output probability (ranging from 0 to 1) and the true trial label (0 for left, 1 for right). Lower DTT values indicate closer alignment between neural representations and the attended side (**Figure 1F**).

### Correlation between decoding performance and behavior

To examine whether neural decoding of spatial attention predicts behavioral performance, we assessed across-participant correlations between decoding accuracy and behavioral measures (reaction time and response accuracy) for each task. Decoding accuracy values were averaged within three 100-ms time windows of interest: a pre-target window (−125 to −25 ms) associated with attention orientation just prior to target presentation, a window around the decoding peak (150–250 ms) and associated with target attentional processing, and a late window (350–450 ms) associated with attention reorienting-related activity that is the case for invalid trials.

For each participant, task, and time window, we computed differences in decoding and behavioral performance relative to the *100 Valid* task, which served as the reference condition (ΔDecoding = *100 Valid* − *90 Valid*; ΔBehavior = *100 Valid* − *90 Valid*). This normalization captured individual deviations in neural and behavioral performance from the baseline condition in which cue validity was perfect. Behavioral measures were taken from both valid and invalid trials of the *90 Valid-IC* and *90 Valid-CC* tasks.

To avoid circularity, decoding models were trained on all trials and tested within subsets defined by task difficulty (easy vs. difficult) and congruency (CS vs. IS). There was no redundance between training and testing trials. Because behavioral accuracy varies most meaningfully on difficult trials, while reaction times (RTs) are more reliable on easy trials, correlations between decoding and accuracy were computed for difficult trials, and correlations between decoding and RT were computed for easy trials. Correlations were calculated separately for IS and CS trials. However, as IS trials more directly reflect attention to the target rather than the distractor, results from IS conditions are reported in the main text, while analyses for CS trials are presented in the Supplementary Information as a control (**Figure S4**).

For each combination of task (*100 Valid, 90 Valid-IC, 90 Valid-CC*), trial validity (valid, invalid), and time window, Pearson’s correlations were computed across participants between ΔDecoding and ΔBehavior between the pair of tasks being considered. The correlation coefficient (*r*) and its corresponding *p*-value were taken directly from the correlation test output. For valid trials, stronger decoding differences (reflecting greater attentional bias) were expected to correlate with slower RTs and lower accuracy; for invalid trials, the opposite relationship was predicted, reflecting more efficient reorienting from the cued to the uncued location. Bonferroni correction was applied within each behavioral endpoint (RT or accuracy) to account for multiple comparisons across tasks and time windows (α = 0.0083).

### Time-Frequency Analysis of DTT

To investigate potential oscillatory components underlying attentional dynamics, we computed time–frequency representations (TFRs) of the decoding-derived DTT time series for each single trial, subject, and task using the Superlet transform (SLT) (Moca et al., 2021). The analysis covered a frequency range of 3–40 Hz, with SLT parameters set to base cycles = 1, minimum order = 3, and maximum order = 20. TFRs were computed over a –900 to +300 ms interval relative to target onset, with an extended pre-stimulus window included to minimize edge effects.

For each subject and condition, we obtained the median TFR across trials and applied a spectral parametrization using the “Fitting Oscillations & One-Over-F” or FOOOF algorithm (Donoghue et al., 2020) to separate periodic from aperiodic (1/f) components, retaining the periodic power spectrum for subsequent analysis. The same procedure was performed for both the attention decoding DTT and a control DTT (decoding distractor difficulty), allowing us to isolate oscillatory components specifically linked to attentional processes.

We computed the log-transformed contrast between attention and control TFRs (log[Att] − log[Ctrl]) for each subject to quantify condition-specific changes in oscillatory power. Group-level statistical testing was performed using a non-parametric cluster-based permutation test (Maris and Oostenveld, 2007) as implemented in MNE-Python. The test used a one-tailed permutation_cluster_1samp_test (tail = 1) with a threshold-free cluster enhancement (TFCE; start = 0, step = 0.2), 1000 permutations, and an adjacency matrix defining neighboring time–frequency points. Significant clusters were identified based on corrected *p*-values (Att > Ctrl).

Group-level TFRs were visualized by computing the median of individual log-contrast TFRs across subjects and highlighting clusters that reached statistical significance. To quantify alpha-band effects, we extracted median power within the late pre-target interval (−175 to −25 ms) and averaged over the 10–12 Hz frequency range for each participant. These values were visualized using half-violin plots, with individual data points shown as dots and the group median indicated by a solid horizontal line.

For statistical comparisons across tasks, we performed a Friedman test on the within-subject alpha-band log-contrast values, followed by two-tailed Wilcoxon signed-rank tests for post hoc pairwise comparisons. Bonferroni correction was applied to adjust for three task comparisons (α = 0.0167). All analyses were performed in log space; dB visualizations correspond to 10× the same log-contrast values and do not affect the underlying statistics.

### Statistics and software

All reported summary values are medians ± standard error of the median (s.e.m.), and all statistical tests were non-parametric due to non-normal data distributions. Pairwise comparisons were evaluated using Wilcoxon signed-rank tests, and omnibus effects were assessed using Friedman’s tests. Bonferroni correction was applied for post hoc multiple comparisons unless otherwise specified.

MEG preprocessing and alpha lateralization visualizations were performed in MATLAB R2023a (The MathWorks, Natick, MA) using the FieldTrip toolbox (release 20211102).

Decoding analyses (both static and time-resolved) and DTT distribution analyses were conducted in MATLAB R2023a. Statistical significance for decoding was evaluated using permutation tests (1,000 iterations with randomly shuffled trial labels) to estimate chance-level performance and generate 95% confidence intervals. For time-resolved decoding, repeated non-parametric tests were performed across time points and corrected using the Benjamini–Hochberg false discovery rate (FDR) procedure, which is more appropriate for partially dependent samples (arising from overlapping sliding windows) than Bonferroni correction.

Time–frequency analyses of the DTT time series were performed in Python 3.12.2. For each subject and task, superlet-based time–frequency representations (TFRs) were computed as periodic (aperiodic-removed) log-power matrices (frequency × time) for attention and control trials. The group contrast was defined per bin as the median of per-subject differences (median of [attention − control]). Statistical inference tested the paired, one-tailed effect (Attention > Control) within –475 to –25 ms using TFCE-based cluster permutation in MNE-Python v1.2.3 (permutation_cluster_1samp_test) with 1000 permutations, TFCE start = 0, step = 0.2, 2-D frequency×time adjacency (combine_adjacency), seed = 42, and cluster-level α = 0.05 (FWER).

## Results

Participants performed a covert spatial attention task while undergoing MEG recordings (**Figure 1**). On each trial, two oriented gratings (target and distractor) appeared simultaneously to the left and right of central fixation in the lower visual field, with each positioned 3° to the left or right of central fixation, respectively. Participants discriminated the orientation of the grating on the cued side (target) relative to an imaginary diagonal reference, indicating whether it deviated clockwise or counterclockwise. To manipulate attentional demands, three cue validity tasks were implemented: a fully valid task in which targets always appeared on the initially cued side (100 Valid), and two 90% valid tasks. In the 90 Valid-IC task (for ipsi-cue 90% Validity task), 90% of trials were valid and invalid trials were signaled by the black square on the attended stimulus flipping position, requiring participants to shift their attention to the opposite side when the cue changed. In this task, the optimal strategy consisted in monitoring the cued grating. In contrast, in the 90 Valid-CC task (for contra-cue 90% Validity task), invalid trials were indicated by a cue flip on the unattended stimulus, requiring participants to monitor the unattended side throughout the block for potential switches. In this task, the optimal strategy consisted in monitoring both the cued grating (to track valid trials) and the uncued grating (to track invalid trials). Eye movements were continuously monitored to ensure central fixation throughout the task. For all analyses, only trials in which no saccades occurred (exceeding 1.5° from central fixation) in the 500 ms preceding target onset and no blinks were detected in this interval were included. Below, we report the results obtained from these analyses.

### Behavioral performance varied as a function of task

Before examining neural decoding, we first verified that manipulations of cue validity and switching rule effectively modulated behavioral performance, confirming that participants adapted their attentional strategy to each task’s demands. Reaction times and accuracy were therefore analyzed to assess how task structure influenced perceptual performance and attentional control when target was presented at the cued location. Reaction times (RTs) were first examined for valid trials across tasks (**Figure 2A**). A repeated-measures ANOVA revealed significant main effects of Task (F(2, 58) = 17.33, p = 1.26 × 10⁻⁶) and Difficulty (F(1, 29) = 28.07, p = 1.11 × 10⁻⁵), indicating slower responses in the *90 Valid* tasks and for difficult trials. No main effect of Congruency was observed (p = 0.153). No interactions reached significance (all p > 0.27). Post hoc Wilcoxon signed-rank tests confirmed that RTs were longer for difficult than easy trials across all tasks (all p ≤ 0.0001). Small congruency effects were found for easy trials in the *100 Valid* (p = 0.0098) and *90 Valid–CC* (p = 0.0387) tasks, with IS trials slightly slower than CS trials. Overall, increased perceptual difficulty reliably slowed behavioral responses under all cue-validity conditions, whereas stimulus congruency had only a minor influence on RTs during valid trials, indicating that subjects were correctly performing the task and were not driven in their response by the uncued grating change.

**Figure 2.**
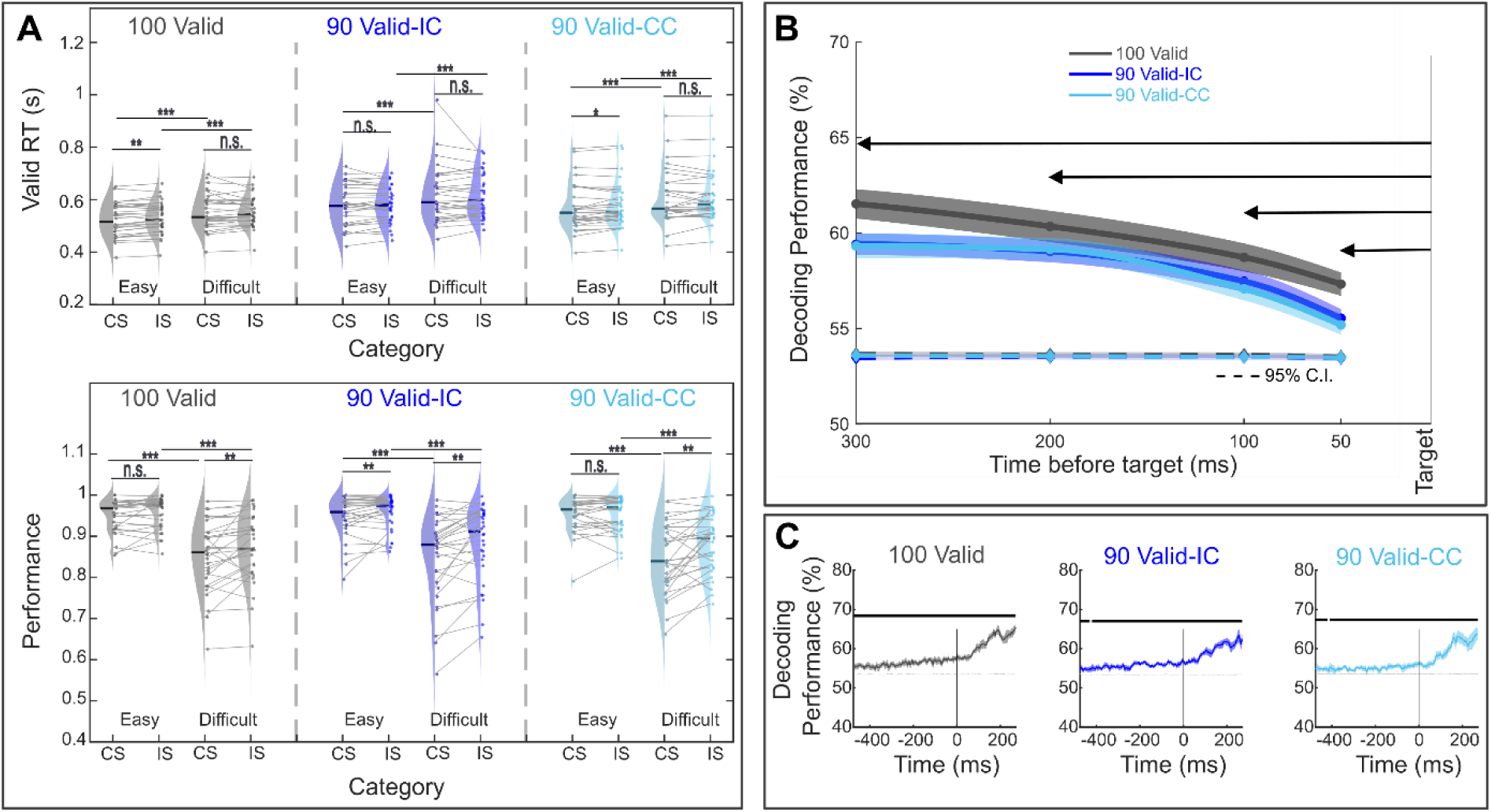
(A) Behavioral results across tasks. Median reaction times (RT; top) and performance accuracy (bottom) for valid trials across three cue-validity tasks: **100 Valid**, **90 Valid-IC**, and **90 Valid-CC.** Within each task, data are shown separately for **Easy** and **Difficult** conditions and for **Congruent (CS)** and **Incongruent (IS)** stimulus categories. Violin plots depict the distribution of participant medians; **solid horizontal lines** indicate group medians, **dots** represent individual participants, and **gray connector lines** link values from the same participant across conditions. Paired Wilcoxon *signed-rank tests* were used for statistical comparisons (*p* < 0.05 (*), *p < 0.01* (**), *p* **< 0.001** (***), n.s. = not significant). **(B) Static decoding for different pre-stimulus window lengths.** Mean neural activity was computed for each MEG visual channel over four pre-stimulus windows of 50 ms, 100 ms, 200 ms, and 300 ms before stimulus onset. For each participant and each time window, an LDA classifier was trained to decode the cued side using these mean amplitudes as features. Decoding performance was calculated separately for each window length and task condition (100 Valid, 90 Valid-IC, 90 Valid-CC). Statistical significance was assessed using permutation tests (1,000 iterations with randomly shuffled trial labels) to estimate chance levels and generate 95% confidence intervals. **(C) Time-resolved decoding of visual channels group-level results. Left panel.** Time-resolved decoding accuracy for the 100 Valid task. The gray solid line shows the median (+/- s.e.) decoding performance across participants at each time point relative to stimulus onset (0 ms). The gray line indicates the median of the 95% confidence intervals across subjects. Solid black horizontal line segments mark time intervals where decoding performance was significantly above chance level (Wilcoxon signed-rank tests with Bonferroni correction). The vertical black dashed line at 0 ms denotes the time of target presentation. **Middel panel.** Same as left panel, for the 90 Valid-IC task. **Right panel.** Same as left panel, for the 90 Valid-CC task.

A 3-factor RM-ANOVA on proportion correct with factors Task (100 Valid, 90 Valid-IC, 90 Valid-CC) × Difficulty (Easy, Difficult) × Congruency (CS, IS) revealed robust main effects of Difficulty (F≈73.99, p≈1.8×10⁻⁹) and Congruency (F≈12.42, p≈0.0014). The Difficulty × Congruency interaction was significant (F≈14.34, p≈7.1×10⁻⁴), indicating that the congruency effect was stronger on Difficult than Easy trials. The Task main effect and Task × Difficulty and Task × Congruency interactions were not significant, but the three-way Task × Difficulty × Congruency interaction reached significance with Greenhouse–Geisser correction (p≈0.047), motivating task-wise follow-ups. Post-hoc tests within tasks showed clear congruency effects on 90 Valid-IC (Easy: M = 0.012, SD = 0.026, p = 0.0201; Difficult: M = 0.036, SD = 0.051, p = 0.0005) and on 90 Valid-CC for Difficult trials (M = 0.036, SD = 0.055, p = 0.0012), but not for Easy trials (M = 0.001, SD = 0.023, p = 0.7483). Consistent Wilcoxon signed-rank tests confirmed lower accuracy on Difficult vs Easy trials within each task (all p < 0.001) and a reliable CS–IS difference on Difficult trials (all p ≤ 0.0013), with smaller or absent CS–IS differences on Easy trials (significant only for 90 Valid-IC, p = 0.0034). In summary, valid-trial accuracy decreases with difficulty, and congruency costs are most evident on difficult conditions, especially in the 90 Valid-IC and 90 Valid-CC tasks.

Overall, these results confirmed that subjects’ reaction time and behavioral accuracies were impacted by task design. Behavioral performance for invalidly cued targets are presented in a later section.

### Spatial attention can be decoded from the MEG sensor-level signals

We first tested whether spatial attention could be reliably decoded from temporal MEG activity patterns extracted from the visual occipital and parietal sensors. Decoding from non-visual or all sensors is occasionally reported for comparison. Using Linear Discriminant Analysis (LDA), classifiers were trained to discriminate whether attention was directed to the left or right grating on each trial. We first examined static decoding performance across increasing pre-stimulus integration windows to determine the temporal scale over which attention-related signals are optimally expressed in the MEG data. We then extended this approach to a time-resolved decoding analysis to capture the moment-to-moment evolution of attentional allocation. By applying a sliding-window classifier across the pre-target interval, we could track how the discriminability of left-versus right-attended trials fluctuated over time, thereby revealing the temporal dynamics and rhythmic structure of the attentional code preceding stimulus onset.

#### Static decoding

Decoding accuracy was computed within pre-stimulus windows ranging from 50 to 300 ms (**Figure 2B**). Across all tasks, accuracy was significantly above chance (all p < 0.0125; Bonferroni-corrected α = 0.0125, permutation-based Wilcoxon tests). A Friedman test confirmed a strong effect of window length on decoding accuracy (p < 1 × 10⁻⁴). Post hoc one-tailed Wilcoxon signed-rank tests (Bonferroni-corrected α = 0.0083) revealed significant increases from 50 ms to 100 ms (p < 1 × 10⁻⁴), 50 ms to 200 ms (p < 1 × 10⁻⁴), 100 ms to 200 ms (p < 1 × 10⁻⁴), 50 ms to 300 ms (p < 1 × 10⁻⁴), and 100 ms to 300 ms (p < 1 × 10⁻⁴), whereas the increment from 200 ms to 300 ms did not reach significance (p = 0.4284). Thus, decoding accuracy increased with longer integration windows, plateauing beyond 200 ms. Taken together, these results mirroring the intracortical attention decoding studies (Astrand et al., 2016; Gaillard et al., 2020), and demonstrate that spatial attention can be robustly decoded from visual sensor-level MEG activity, with decoding strength increasing for longer pre-target integration windows and remaining consistently above chance across all tasks.

#### Time-resolved decoding

A complementary group-level time-resolved analysis (**Figure 2C**) using 50 ms sliding Hann-weighted windows (5 ms steps) revealed that decoding accuracy dynamically evolved throughout the pre-target period. Permutation testing (1,000 label shuffles) was used to derive subject-specific 95% confidence intervals, and intervals of significant above-chance decoding were identified via Wilcoxon signed-rank tests with Benjamini–Hochberg correction. Solid black lines on the traces indicate time points where decoding accuracy exceeded chance. As can be seen in **Figure 2C**, at the group level, time-resolved decoding was significant across all time points from pre-cue to post cue interval, to the exception of a few time points in the very early pre-target interval in the *90 Valid* tasks). Importantly, single-subject examples (**Figure 3A, supplementary Figure S2**) illustrate consistent above-chance decoding across all task conditions for most subjects, possibly indicating a correlation between decoding accuracy and behavioral performance. This will be addressed in the next sections. When all channels were used for the decoding, a similar result as reported in **Figure 2C** is observed (**Figure S3**, bottom panel). Exclusively considering non-visual channels deteriorated decoding significance, mostly in the pre-target epoch (**Figure S3**, top panel). Together, these results demonstrate that the spatial attention orientation can be reliably decoded from human MEG signals with high temporal precision, primarily driven by activity in visual sensors, and that this decoding remains robust across individuals and task variants.

**Figure 3.**
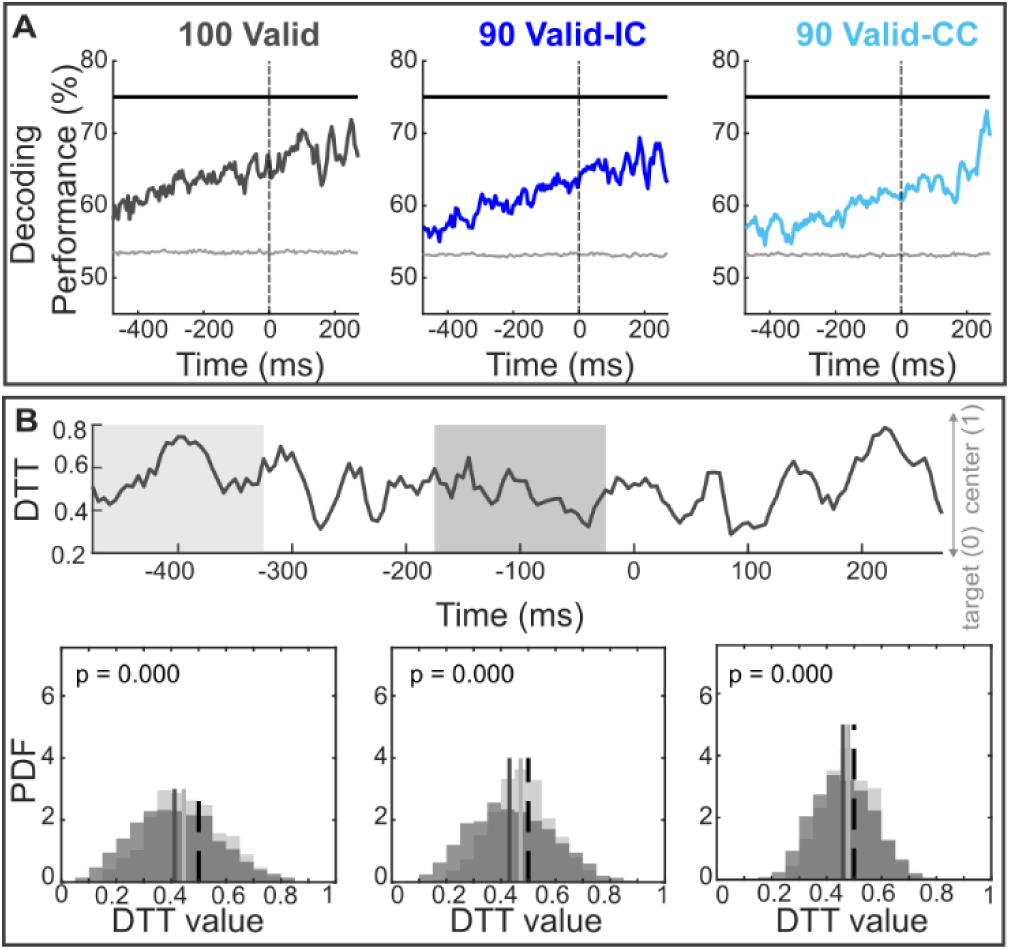
Time-resolved decoding and DTT distributions for an example subject. **(A)** Decoding performance over time for each of the three tasks for subject 126. All as in figure 2C. **(B)** Time series of average DTT values across all trials for each task. The shaded light gray area indicates the early analysis window (–475 ms to –325 ms), and the shaded dark gray area marks the late window (–175 ms to –25 ms), immediately preceding target onset. Lower DTT values reflect classifier outputs closer to the true attended side, indicating more precise attentional allocation. **(C)** Probability density distributions of DTT values within the early (light gray) and late (dark gray) windows for each task. The dashed vertical line at 0.5 indicates a DTT equidistant to both left and right cued locations.

### Reduced cue validity weakens neural discriminability

Having established that spatial attention can be robustly decoded across tasks, we next investigated whether task structure, as defined by cue validity and switching rules, modulates overall decoding strength of the attentional signal. We thus examined differences in decoding accuracy between tasks irrespective of window size. A Friedman test revealed a significant effect of task (χ²(2) = 12.87, p = 0.0016), with median decoding accuracy of 59.88% (IQR = 57.04–61.92%) for the *100 Valid* task, 58.17% (IQR = 55.98–60.21%) for the *90 Valid-IC* task, and 57.85% (IQR = 55.97–60.10%) for the *90 Valid-CC* task. Pairwise Wilcoxon signed-rank tests with Bonferroni correction (α = 0.0167) showed that decoding accuracy was significantly higher for *100 Valid* compared to both *90 Valid-IC* (p = 0.0009) and *90 Valid-CC* (p = 0.0003), whereas the difference between *90 Valid-IC* and *90 Valid-CC* was not significant (p = 0.5857). To investigate task differences within each specific window size (50, 100, 200, and 300 ms before target onset), additional pairwise Wilcoxon signed-rank tests with Bonferroni correction (α = 0.0167) were conducted. Significant reductions in decoding accuracy from *100 Valid* to *90 Valid-IC* were observed at all window sizes (50 ms: p = 0.0171, 100 ms: p = 0.0005, 200 ms: p = 0.0024, 300 ms: p = 0.0014). Similarly, decoding accuracy was significantly lower in *90 Valid-CC* compared to *100 Valid* across all windows (50 ms: p = 0.0026, 100 ms: p = 0.0006, 200 ms: p = 0.0027, 300 ms: p = 0.0029). In contrast, no significant differences were found between *90 Valid-IC* and *90 Valid-CC* at any window size (all p > 0.33). These results demonstrate that reducing cue validity reliably diminishes the strength of the neural attention code, whereas differences arising from rule switching will be uncovered in subsequent analyses.

### Attention is more aligned with the cued target in Late window but effect differs across tasks

Because decoding accuracy provides a binary measure of classifier performance that may obscure trial-by-trial variability in attentional precision as well as within-trial attentional dynamics, we next examined the Distance to Truth (DTT) metric, a continuous index of how closely the classifier output aligns with the true attended location, to capture finer-grained changes in attentional focus over time. We examined how attentional allocation evolved as the target onset approached, and we compared DTT values between an early (−475 to −325 ms) and a late (−175 to −25 ms) pre-target window. **Figure 3B** illustrates, for a representative subject, a single trial DTT trace in time (top panel) as well as the probability distributions of DTT across trials in the early and late pre-target windows. DTT probability distributions were significantly smaller in the late window as compared to the early window, indicating that attention was significantly closer to the cued target in the later window.

This observation was confirmed across subjects. Indeed, at the group level, Wilcoxon signed-rank tests confirmed significant reductions in DTT from the early to late window in the *100 Valid* (p = 0.002) and *90 Valid–IC* (p = 0.017) tasks, thus reflecting a shift of attentional focus toward the target side. This was however not the case for the *90 Valid–CC* task as DTT decrease from the early to late window did not reach significance (p = 0.318; **Figure 4A**).

**Figure 4.**
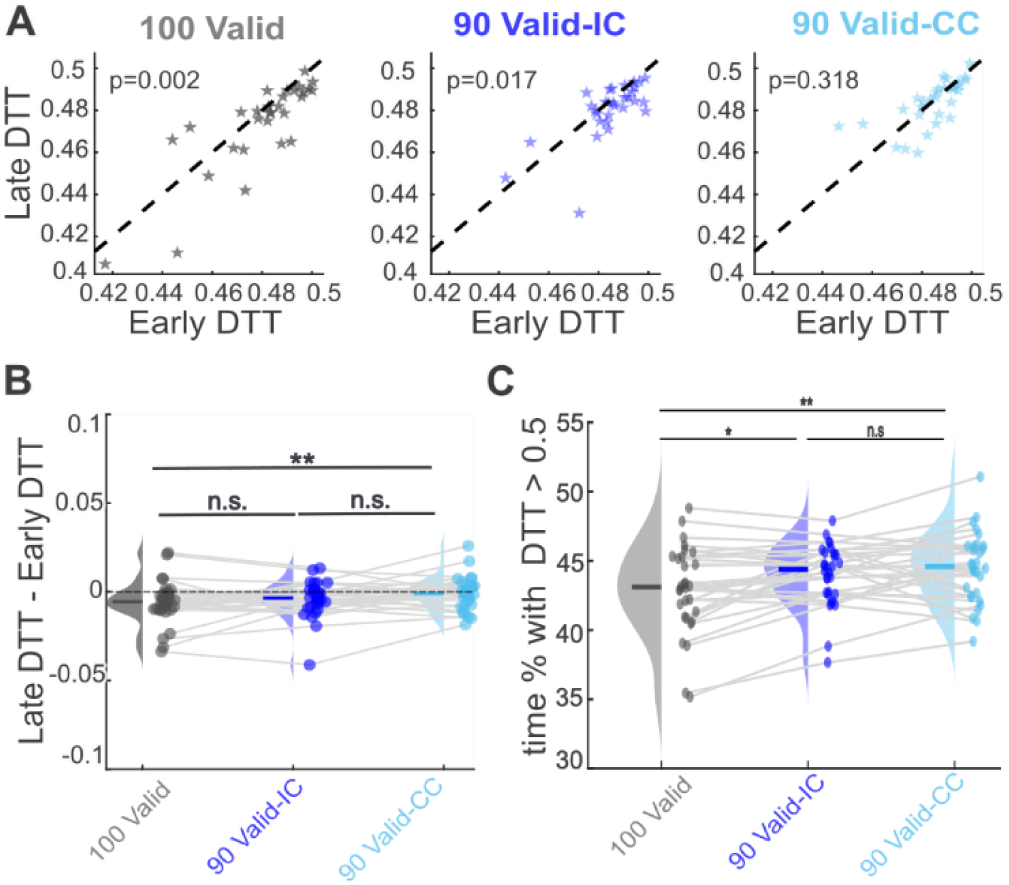
Median DTT comparisons across early and late pre-stimulus intervals. **(A) Left panel**, Scatter plot showing, for each subject, the median DTT in the early window (–475 to –325 ms) versus in the late window (–175 to -25 ms) for the 100 Valid task (p-value, Wilcoxon signed-rank test). Each star represents one participant. The diagonal dashed line indicates equality between early and late medians. Points falling below the diagonal reflect trials where DTT decreased (i.e., attention became more aligned with the cued target) closer to stimulus onset. **Middle panel**, same as left panel, but for the 90 Valid-IC task. **Right panel**, same as left panel, but for the 90 Valid-CC task. **(B)** Distribution plots of the within-subject differences in median DTT between the late and early windows for each task. For each participant, the median DTT in the early window was subtracted from the median DTT in the late window, such that negative values indicate a reduction in DTT over time (i.e., improved attentional alignment). Half violin plots illustrate the distribution of these differences across subjects for each condition. Small dots show individual data points, with connecting lines linking the tasks within each participant. Horizontal colored lines indicate median differences per condition. The dashed line at zero represents no change in DTT. * p < 0.05, ** p < 0.01, *** p < 0.001 (Wilcoxon rank-sum tests with Bonferroni correction for multiple comparisons); “n.s.” denotes no significant difference between conditions. **(C) Comparison of late-window attentional alignment between tasks.** This figure shows the proportion of DTT values exceeding 0.5 during the late pre-stimulus window (–175 to –25 ms) for each subject and task: dark gray (100 Valid); dark blue (90 Valid-IC) and light blue (90 Valid-CC). Individual subject values are represented as dots, with gray connecting lines linking measurements between tasks for each participant. Horizontal colored lines within each half-violin indicate the median proportion per condition. * p < 0.05, ** p < 0.01, *** p < 0.001 (Wilcoxon rank-sum tests with Bonferroni correction for multiple comparisons).

To test whether this temporal change differed across tasks, we computed for each participant the Late–Early DTT difference and compared these values using one-tailed paired Wilcoxon signed-rank tests with Bonferroni correction (α = 0.0167). The decline was significantly stronger in the *100 Valid* task compared to the *90 Valid–CC* task (p = 0.0090, z = −2.37), while differences between *100 Valid* and *90 Valid–IC* (p = 0.1938, z = −0.86) and between *90 Valid–IC* and *90 Valid–CC* (p = 0.0721, z = −1.46) were not significant. A Friedman test across tasks approached significance (χ²(2) = 5.40, p = 0.0672), suggesting a trend toward task-dependent modulation of the DTT decline (**Figure 4B**). Overall, DTT values decreased as the moment of target presentation neared, consistent with a growing attentional bias toward the target location. Although the overall task effect did not reach statistical significance, the observed pattern supports the hypothesis that this attentional bias was strongest under fully valid cueing (100 Valid) and weakened progressively with reduced cue validity (90 Valid–IC, 90 Valid–CC).

### Dwell time contralateral to the cued location varies as a function of the task

To further characterize how task demands influenced the spatial dynamics of attention, we next examined the dwell time of the decoded attentional spotlight, that is, how long attention remained contralateral versus ipsilateral to the cued location during the late pre-target interval. Building on the previous analyses, we quantified the proportion of time points within the late pre-target window (−175 to −25 ms) in which the DTT was positioned contralateral to the cued location—that is, instances where attentional tracking crossed the midline (**Figure 4C**). For each participant and task, this proportion was calculated as the fraction of DTT values exceeding the midline threshold (0.5). Consistent with the task design, we expected more frequent midline crossings in conditions with reduced cue validity. A Friedman test revealed a significant overall task effect (χ²(2) = 13.07, p = 0.00145). Post hoc one-tailed Wilcoxon signed-rank tests (Bonferroni-corrected α = 0.0167) confirmed higher proportions of midline-crossing DTTs in both the *90 Valid–IC* (p = 0.0162, r = 0.391) and *90 Valid–CC* (p = 0.0002, r = 0.642) tasks compared with the 100 *Valid task*, indicating medium and large effects, respectively. The difference between the two 90 Valid tasks was close to significance (p = 0.0666, r = 0.274). These results confirm a reliable increase in midline-crossing DTTs under conditions of reduced cue validity, with the strongest effect observed in the *90 Valid-CC* task. This is in agreement with the difference in task rule switching differences between the *90 Valid-IC* and *90 Valid-CC* tasks. Indeed, in the *90 Valid–CC* task, participants were required to continuously monitor the unattended side for potential cue flips, promoting a more distributed or exploratory attentional strategy and resulting in more frequent midline crossings. In contrast, the *90 Valid–IC* task primarily engaged reactive shifts of attention in response to explicit switch cues, yielding comparatively fewer cross-hemifield transitions, although subjects might have engaged weaker attentional engagement at the cued location in order to promote faster switching reaction times. Together, these findings indicate that when cue reliability decreases and task rules encourage sustained monitoring of multiple spatial locations, the decoded attentional spotlight becomes more labile and less tightly anchored to the cued target position.

### Decoding of spatial attention correlates with behavioral performance

Having established that task structure modulates the strength and spatial dynamics of attentional decoding, we next asked whether variability in neural decoding strength translates into behavioral performance differences. If the fidelity of the decoded attentional signal reflects the precision of attentional deployment, stronger decoding should be associated with faster and more accurate responses. To test this hypothesis, we examined across-participant correlations between decoding accuracy and behavioral measures (reaction time and accuracy) across tasks and time windows.

To assess the link between neural decoding strength and behavior, we examined across-participant correlations between changes in decoding accuracy and behavioral measures, normalized to the *100 Valid* task (i.e. ΔDecoding = *100 Valid − 90 Valid* vs. ΔBehavior = *100 Valid − 90 Valid*). Correlations were computed separately for the *90 Valid–IC* and *90 Valid–CC* tasks. Because easy target trials resulted in only subtle differences in accuracy across tasks, we focused, for these trials, on RTs. In contrast on difficult target trials, differences in accuracy were highly task dependent while reaction times were overall slowed down. As a result, for these difficult trials, we focused on accuracy variations. Last, we focused on IS trials as these unambiguously identified correct trials as compared to CS trials in which the overt response could have been influenced by either the target or the distractor. All non-reported conditions are nonetheless presented in supplementary material. Three 100-ms temporal windows were tested (−125 to −25 ms, corresponding to pre-target epoch; 150–250 ms, corresponding to initial attentional capture by attention; and 350–450 ms, corresponding to putative attentional capture following attentional switches, see **supplementary Figure S4** and associated description), with Bonferroni correction applied across tasks × windows (α = 0.0083). We first focused on valid trials, where decoding strength directly indexes the efficiency of sustained attention toward the cued target, before turning to invalid trials, which probe how attentional bias and flexibility influence reorienting performance under changing task demands.

#### Behavioral relevance of attentional decoding under valid cueing

We examined, in valid trials, whether fluctuations in attention decoding strength predicted individual differences in behavioral accuracy and reaction time across tasks. We first focused on difficult IS trials (**Figure 5A**, corresponding analyses for CS trials are presented in **supplementary Figure S5**). A significant positive correlation was observed between differences in behavioral accuracy between the *100 Valid* and the *90 Valid–CC* task during the pre-target (−125 to −25 ms, p = 0.0018) and between the *100 Valid* and the *90 Valid–IC* task during the pre-target (150–250 ms, p = 0.0045) windows, indicating that greater reductions in decoding were associated with larger accuracy losses relative to the 100 Valid baseline (**Figure 5A**). No other task–window combination reached significance (all p ≥ 0.01). We next focused on easy IS trials (**Figure 5B**). No significant correlations were observed after correction (all p ≥ 0.01). Thus, overall, during valid trials, reduced attentional decoding in the 90 Valid tasks reliably tracked decreases in behavioral accuracy, whereas corresponding RT effects were not significant.

**Figure 5.**
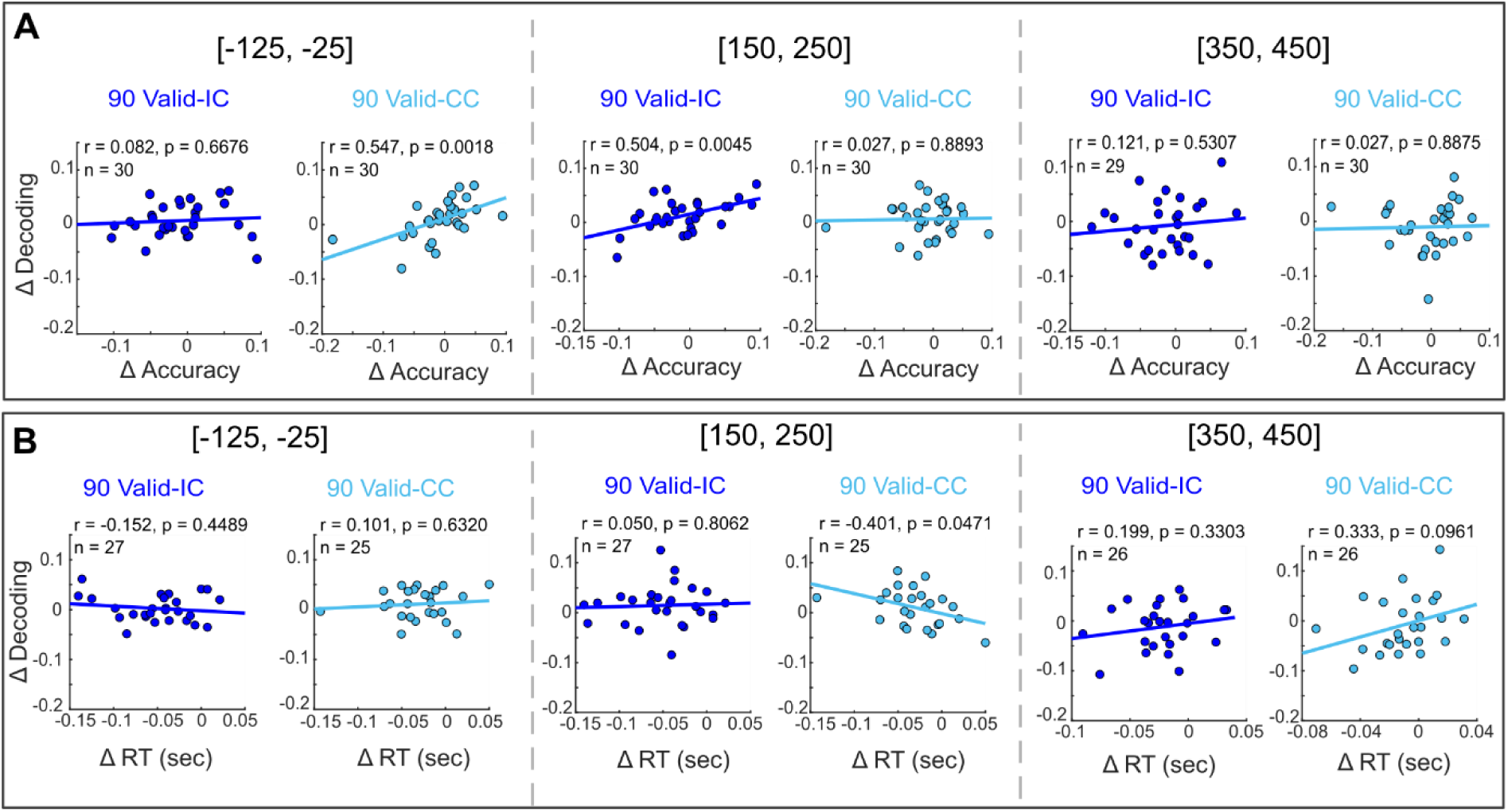
Correlation between decoding and behavioral differences across tasks for valid trials. (A) Correlation between changes in decoding accuracy and behavioral performance (accuracy) across participants for the Difficult–IS condition. (B) Correlation between changes in decoding accuracy and reaction RT for the Easy–IS condition. For each participant, differences were computed relative to the 100 Valid task, using it as a reference for valid trials from the 90 Valid-IC and 90 Valid-CC tasks. Correlations are shown for three temporal decoding windows ([-125, −25], [150, 250], [350, 450] ms). Each dot represents one participant; solid lines denote least-squares linear fits. Pearson’s *r*, *p*-values, and sample sizes (*n*) are reported within each panel. Participants with accuracy below 50% or RTs exceeding ± 2 s.d. of the median in valid trials were excluded from the respective analyses.

#### Behavioral relevance of attentional decoding under invalid cueing and switch instruction

We next investigated whether the relationship between decoding strength and behavior extended to invalid (switch) trials, which required participants to reorient attention in response to a cue change. These trials provide a complementary test of attentional flexibility, probing whether stronger neural bias toward the unattended side facilitates faster and more accurate reorienting when the target unexpectedly appears there.

Behavioral performance during attentional reorienting is shown in **Figure 6A**. Median reaction times (top) and performance accuracies (bottom) are displayed for the two 90 % validity tasks (*90 Valid–IC* and *90 Valid–CC*), separately for easy and difficult trials and for congruent (CS) versus incongruent (IS) stimulus categories. Reaction times (RTs) were significantly slower and accuracies lower in invalid (**Figure 6A**) compared with valid trials (**Figure 2A**) for both 90 Valid tasks, confirming a strong cue-invalidity effect (*90 Valid–IC*: RT z = 4.75, p < 0.0001; Accuracy z = –4.71, p < 0.0001; *90 Valid–CC*: RT z = 4.73, p < 0.0001; Accuracy z = –4.77, p < 0.0001). The median RT increase between valid and invalid trials was 0.38 s in the *90 Valid–IC* task and 0.30 s in the *90 Valid–CC* task, accompanied by accuracy drops of 7.6 % and 22 %, respectively. These observations thus confirmed a strong cue invalidity effect. particularly for incongruent trials, suggesting increased demands on attentional monitoring and conflict resolution. In contrast, no reliable RT difference was observed between the two 90 Valid tasks for either valid (z = –0.37, p = 0.64) or invalid (z = –1.46, p = 0.93) trials, indicating comparable response latencies across cueing contexts. Accuracy, however, was significantly lower in the *90 Valid–CC* task relative to the *90 Valid–IC* task for invalid trials (z = –4.42, p < 0.0001, **Figure 6A**), while no difference emerged for valid trials (z = –0.62, p = 0.27). These results demonstrate that cue invalidity robustly impairs both speed and accuracy, and that the cost is especially pronounced in the *90 Valid–CC* task, where monitoring the unattended side imposes higher demands on attentional control. Accuracy was higher for CS than IS trials in the *90 Valid-IC* condition (p < 0.0001), suggesting that participants occasionally missed the cue on the unattended side but still responded correctly due to stimulus congruency. When taking into account target difficult and target/distractor congruency, the ANOVA showed significant main effects of Task (F(1, 28) = 37.67, p = 1.09 × 10⁻⁶), Difficulty (F(1, 28) = 33.62, p = 2.77 × 10⁻⁶), and Congruency (F(1, 28) = 28.01, p = 1.13 × 10⁻⁵), along with Task × Difficulty (F(1, 28) = 9.46, p = 0.0045) and Task × Congruency (F(1, 28) = 67.70, p = 4.52 × 10⁻⁹) interactions. Follow-up Wilcoxon tests indicated no significant differences in RTs between easy and difficult conditions (all p > 0.16). However, strong congruency effects emerged in the *90 Valid–CC* task, where IS trials were slower than CS trials for both easy (p < 0.0001) and difficult (p = 0.0001) conditions. This probably reflects the fact that CS invalid trials aggregate both switch and non-switch trials. No congruency effects were found in the *90 Valid–IC task* (all p > 0.46).

**Figure 6.**
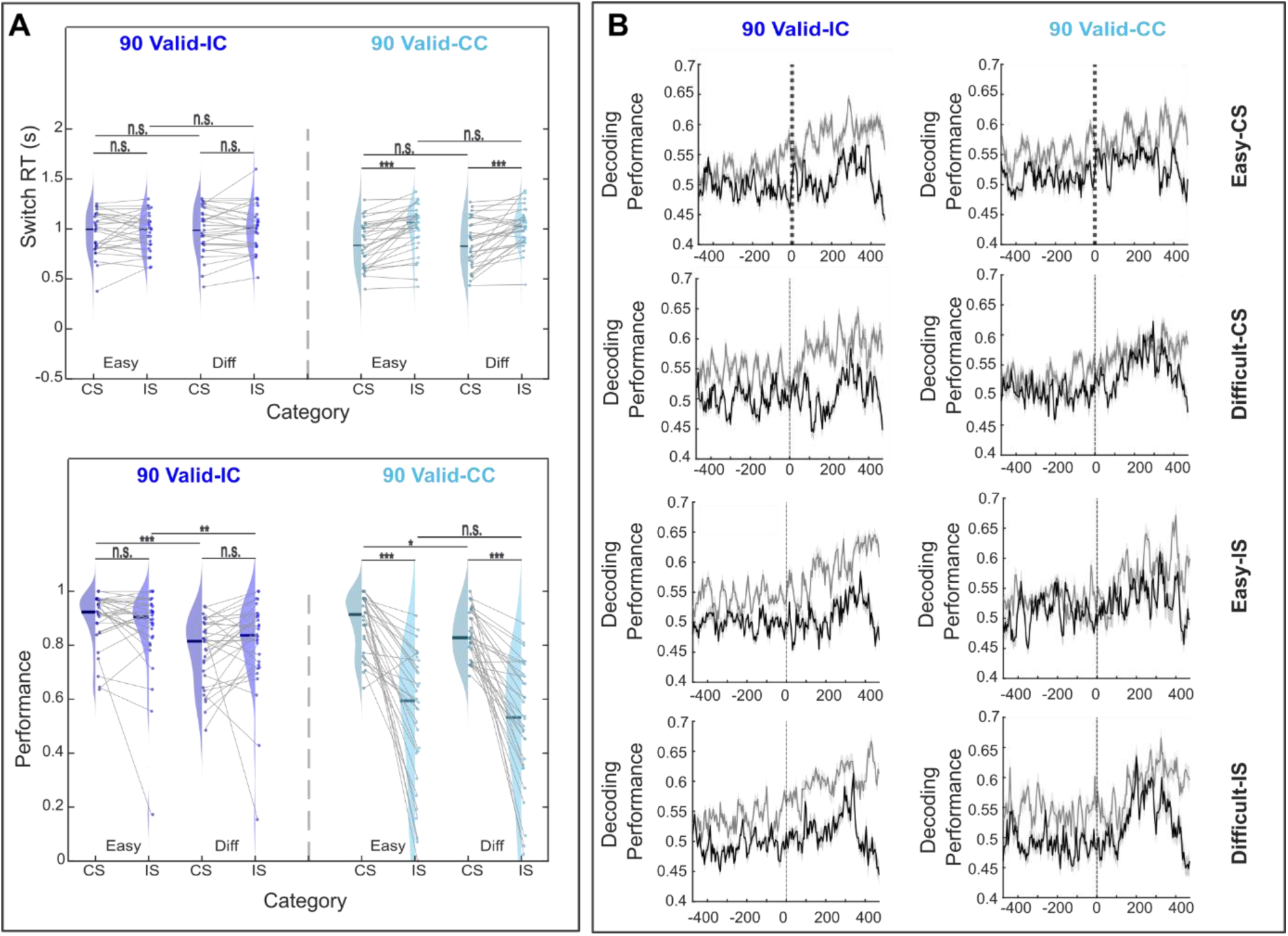
(A) Behavioral results across 90 validity tasks. Median reaction times (RT; top) and performance accuracy (bottom) for Invalid trials across 90 Valid-IC, and 90 Valid-CC tasks. Within each task, data are shown separately for Easy and Difficult conditions and for **CS** and **IS** stimulus categories. Violin plots depict the distribution of participant medians; solid horizontal lines indicate group medians, dots represent individual participants, and gray connector lines link values from the same participant across conditions. Paired Wilcoxon *signed-rank tests* were used for statistical comparisons (*p* < 0.05 (*), *p < 0.01* (**), *p* < 0.001 (***), n.s. = not significant). **(B)** Decoding performance for valid (gray line) vs Invalid (black line) trials for 90 Valid-IC (left panel) and 90 Valid-CC (right panel) tasks across four categories of Easy-CS (first row), Difficult-CS (second row), Easy-IS (third row), and Difficult-IS (last row).

Decoding performance during attentional reorienting is shown in Figure 6B. Given the behavioral evidence for slower and less accurate responses during invalid trials, particularly in the *90 Valid–CC* task, we next asked whether these performance costs reflect changes in the underlying neural representation of spatial attention. Specifically, we tested whether the attentional code established during valid trials remains stable or reorganized when attention must be reoriented. To this end, trials were first divided into four categories according to task difficulty (easy or difficult) and stimulus congruency (CS or IS). For each category, a decoder was trained on valid trials only and subsequently tested on both valid and invalid trials of the same category. The resulting time-resolved decoding traces are displayed in **Figure 6B**, with gray lines indicating decoding performance for valid trials and black lines for invalid trials. Across most conditions, decoding accuracy was reliably lower for invalid than for valid trials, suggesting that attentional representations became less distinct when participants anticipated possible cue switches, likely reflecting the formation of probabilistic expectations about switch timing within each block.

Having established that decoding performance decreased during invalid trials, we next asked whether this neural attenuation was behaviorally meaningful, that is, whether fluctuations in decoding strength predicted variability in reaction times and accuracy across participants. We examined correlations between decoding differences and behavioral performance (reaction time and accuracy) across tasks and time windows as we did, in the previous section, for valid trials. For accuracy (difficult–IS), task-specific patterns emerged in line with cue position (**Figure 7A**, corresponding analyses for CS trials are presented in **supplementary Figure S5**). In the *90 Valid–IC* task (cue on the attended side), positive trends appeared in the mid-latency (150–250 ms, p = 0.0241) windows, consistent with the prediction that stronger decoding differences (weaker cue monitoring) lead to poorer accuracy. In contrast, in the *90 Valid–CC* task (cue on the unattended side), a weak positive correlation in the early window (p = 0.0177) did not match the expected negative direction. None of the accuracy effects survived Bonferroni correction (α = 0.0083).

**Figure 7.**
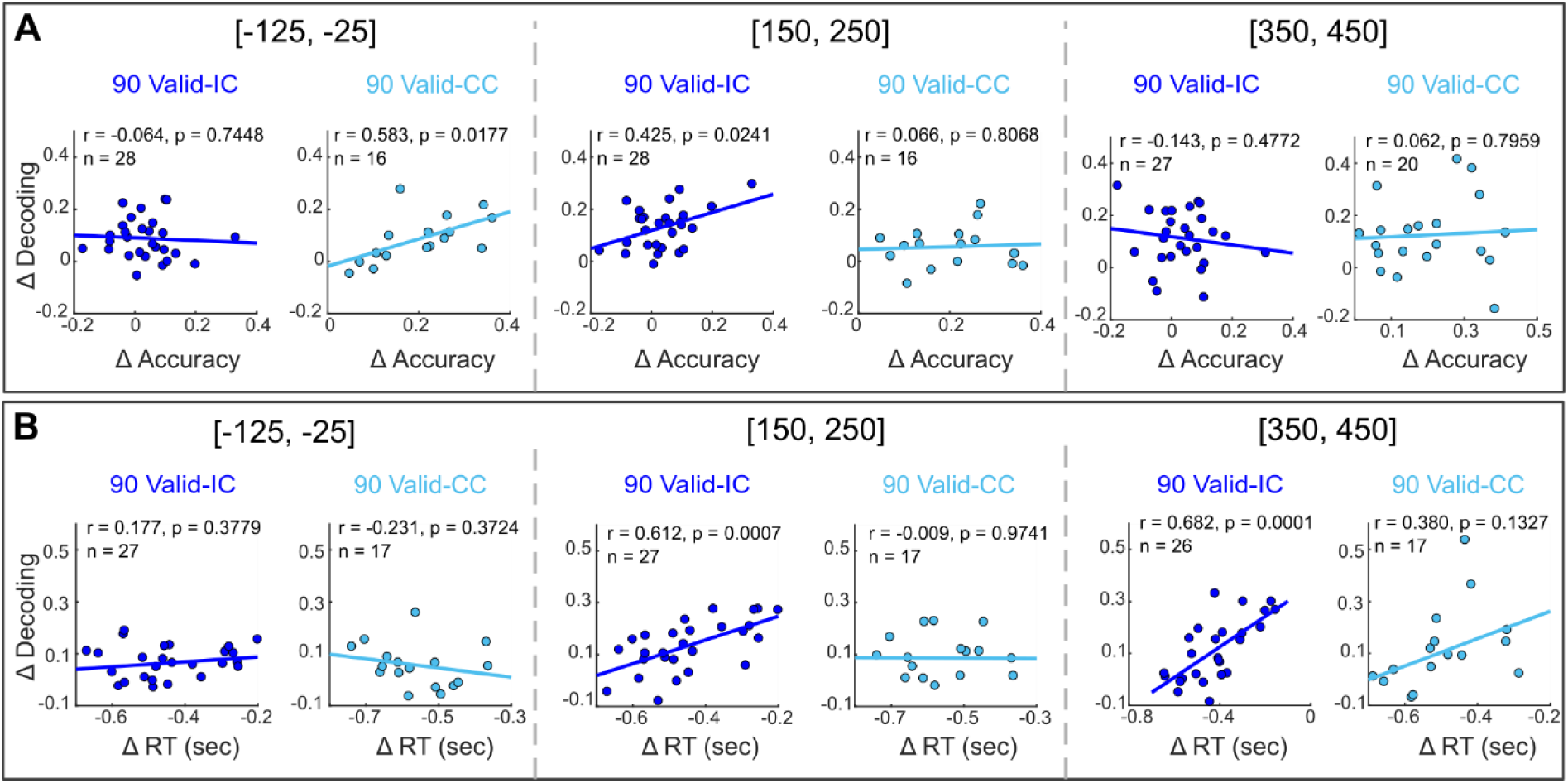
Correlation between decoding and behavioral differences across tasks for invalid trials. (A) Correlation between changes in decoding accuracy and behavioral performance (accuracy) across participants for the Difficult–IS condition. (B) Correlation between changes in decoding accuracy and reaction RT for the Easy–IS condition. For each participant, differences were computed relative to the 100 Valid task, using it as a reference for invalid trials from the 90 Valid-IC and 90 Valid-CC tasks. Correlations are shown for three temporal decoding windows ([-125, −25], [150, 250], [350, 450] ms). Each dot represents one participant; solid lines denote least-squares linear fits. Pearson’s *r*, *p*-values, and sample sizes (*n*) are reported within each panel. Participants with accuracy below 50% or RTs exceeding ±2 s.d. of the median in valid trials were excluded from the respective analyses.

A positive relationship was predicted between ΔDecoding and ΔRT: when decoding differences were large (stronger bias toward the unattended side), participants were faster to respond once the target appeared on the opposite side, yielding smaller ΔRTs. Consistent with this hypothesis, significant positive correlations were found in the *90 Valid–IC* task during the mid-latency (150–250 ms, p = 0.0007) and late (350–450 ms, p = 0.0001) windows, both surviving Bonferroni correction (**Figure 7B**). No significant RT correlations were observed in the *90 Valid–CC* task (all p ≥ 0.01).

Together, these results indicate that neural decoding strength predicts behavioral performance in a task-specific manner: during valid trials, weaker attentional coding was associated with reduced accuracy, particularly in the *90 Valid–CC* condition, whereas during invalid trials, stronger attentional bias toward the unattended side predicted faster reorienting responses in the *90 Valid–IC* condition.

### Rhythmic modulation of the attentional code in the alpha band

The previous sections demonstrate that attentional decoding strength varies with task structure and behavioral performance. We next investigated whether these fluctuations exhibit rhythmic organization, an established hallmark of attentional sampling in both humans and nonhuman primates. To this end, we analyzed the spectral content of the DTT time series to determine whether the decoded attentional signal oscillated at characteristic frequencies. We used Superlet transform (3–40 Hz) and isolated periodic power components with FOOOF. For each subject and task, the logarithmic contrast between attention and control decoding signals (log[Att] − log[Ctrl]) was computed, and statistical significance of condition differences was assessed with a one-tailed cluster-based permutation test (Att > Ctrl, TFCE, 1,000 permutations). At the group level, all tasks exhibited a significant cluster in the alpha frequency range (∼8–12 Hz), indicating a reliable alpha-band modulation of attentional tracking before target onset (**Figure 8A**). To quantify this effect, we extracted median log-contrast power in the 10–12 Hz range within the late pre-target window (−175 to −25 ms). These values were compared across tasks using a Friedman test, which revealed no significant overall difference (χ²(2) = 1.87, p = 0.393). Post hoc Wilcoxon signed-rank tests with Bonferroni correction (α = 0.0167) showed a significant difference between the *90 Valid–IC* and *90 Valid–CC* tasks (p = 0.0164), whereas the contrasts between *100 Valid* and *90 Valid–IC* (p = 0.6263) and between *100 Valid* and *90 Valid–CC* (p = 0.0497) did not reach significant after correction. Together, these results reveal a robust alpha-band modulation of the attentional decoding signal preceding target onset, with subtle task-dependent differences emerging between the two 90 Valid conditions.

**Figure 8.**
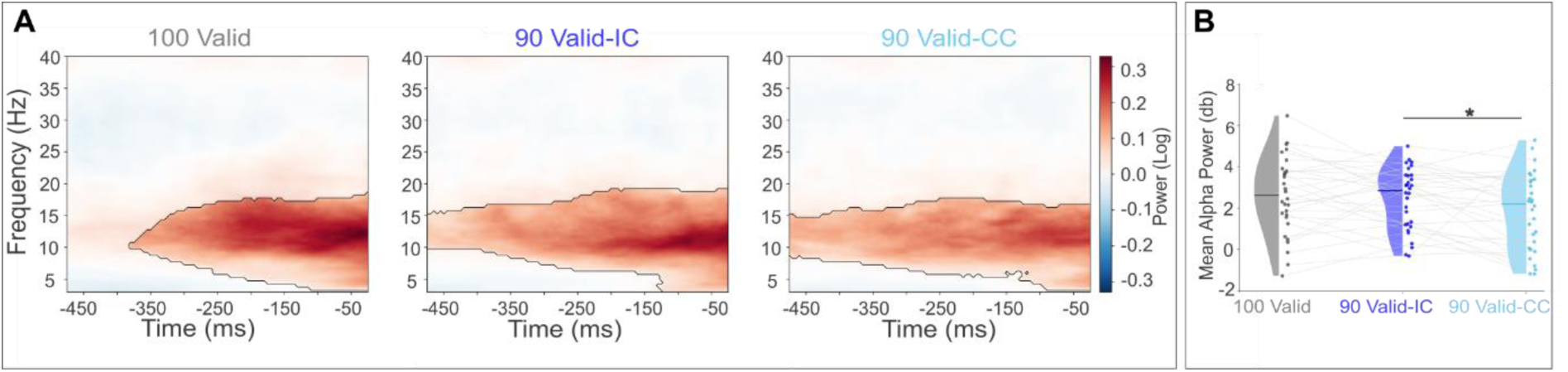
Time-frequency representation (TFR) of DTT time series visual channels. **(A)** Left panel, Contrast TFR for the 100 Valid task, showing significant clusters (with non-significant areas displayed with reduced transparency). Middle panel, Contrast TFR for the 90 Valid-IC task, with significant clusters. Right Panel, Contrast TFR for the 90 Valid-CC task, with significant clusters. **(B)** Median alpha power (10–12 Hz) in the late pre-stimulus window (–175 to –25 ms). Dots indicate individual subject medians, and solid lines within each half-violin show the median across subjects. * p < 0.05, ** p < 0.01, *** p < 0.001 (Wilcoxon rank-sum tests with Bonferroni correction for multiple comparisons).

Our prediction was that alpha band power in the DTT trace should be highest in the 90 Valid tasks that involve switching, and more so in the *90 Valid-CC* task that involves monitoring of both cued and uncued sides. The experimental observations were actually opposite to this prediction. Remarkably, when this analysis was performed using the non-visual channels, significant alpha band DTT trace modulation was only observed for the 90 Valid CC tasks (**supplementary Figure 6A**). However, no clear statistical difference could be seen between the three tasks in the median log-contrast power in the 10–12 Hz range within the late pre-target window (−175 to −25 ms, **supplementary Figure 6B**). When considering all channels, the DTT power in the alpha band was dominated by the pattern observed in when considering just the visual channels though no statistically significant difference could be identified between the three tasks (**supplementary Figure 7**).

## Discussion

The present study establishes that covert spatial attention can be *non-invasively decoded* from human MEG recordings with millisecond precision. Using linear classifiers applied to sensor-level signals, we could reconstruct, on a single-trial basis, the spatial locus of attention throughout the pre-target interval. This demonstrates that the large-scale neural dynamics captured by MEG contain sufficient information to track the moment-to-moment deployment of attention, extending prior EEG and MEG decoding studies (Bae and Luck, 2018; Foster et al., 2017; Samaha et al., 2016; Wyart et al., 2015) by showing that attentional spatial orientation information can be recovered robustly even when task contingencies vary. Decoding accuracy systematically decreased with decreased cue reliability, confirming that the strength and spatial precision of attentional engagement depend on predictive context. The Distance-to-Truth analysis further revealed that attentional focus sharpened as target onset approached, but that this pre-target alignment weakened when cue validity was reduced, indicating more diffuse or exploratory allocation of attention under uncertainty. Task-related differences in decoding strength were mirrored by behavioral performance: stronger attentional decoding across tasks predicted higher differences in accuracy and responses, establishing a tight link between neural coding of attention and behavioral efficiency. Finally, spectral analysis of the decoding trajectories uncovered rhythmic fluctuations in the alpha band (∼8–12 Hz) across all task variants, demonstrating that attention samples space rhythmically even when task structure was highly predictive. Together, these findings validate the feasibility of MEG-based decoding of attention and provide a methodological bridge between invasive primate electrophysiology and human whole-brain recordings.

### Sensor-space versus source-space decoding

While our main analyses focused on sensor-level MEG signals, this choice was guided by both methodological and conceptual considerations. Decoding from sensors provides a direct, high-temporal-resolution readout of distributed neural dynamics without requiring model-based inverse solutions, thereby minimizing assumptions about source geometry and spatial priors (Grootswagers et al., 2017; King and Dehaene, 2014). Restricting analyses to visual sensors allowed us to isolate the sensory components most strongly modulated by covert attention, in line with canonical findings of alpha-band lateralization over occipitoparietal areas (Thut et al., 2006; Worden et al., 2000). Decoding accuracies were degraded when considering only non-visual sensors. Combining non-visual and visual sensors did not significantly improve overall decoding accuracies relative to using just visual sensors suggesting that most of the attention-related information is captured by these latter sensors.

However, spatial attention arises from coordinated interactions within a large-scale frontoparietal network (Corbetta and Shulman, 2002; Kastner et Ungerleider, 2000), and sensor-space decoding cannot disentangle the relative contributions of these distributed sources. Moving to source-localized decoding thus constitutes a critical next step. Source reconstruction methods, such as beamforming or minimum-norm estimation, can project sensor signals into anatomically constrained cortical space, allowing more direct mapping between decoding performance and the underlying neural generators (Cichy et al., 2014; Jaiswal et al., 2020; Konttila et al., 2014). Several recent studies have demonstrated that source-space decoding enhances spatial specificity and improves interpretability of attentional dynamics by revealing localized alpha generators in occipital and parietal cortices as well as task-dependent modulations in prefrontal areas (Foster et al., 2017). Integrating such source-resolved decoding with the trajectory-based approach used here would bridge population-level dynamics to anatomically grounded mechanisms, ultimately linking rhythmic attentional sampling to the coordinated activity of specific cortical subnetworks.

### Task-dependent modulation of the decoded attentional code

Extending prior work, we show for the first time that attentional decoding from MEG generalizes across tasks that manipulate cue validity, revealing that task structure—specifically, cue reliability and switching rules—systematically modulates both the strength and spatial distribution of attention. A growing body of studies has highlighted that attentional control signals are highly dynamic in both space and time (Fiebelkorn et al., 2018, 2013; Hamed, 2025; Landau et al., 2015; Landau and Fries, 2012). Our previous invasive research in macaques established real-time decoding of the two-dimensional locus of covert attention (Astrand et al., 2016, 2020) and revealed its rhythmic (Gaillard et al., 2020) and state-dependent dynamics (Amengual et al., 2022). These invasive studies demonstrated how prefrontal networks flexibly implement proactive and reactive mechanisms of visual suppression (Di Bello et al., 2022) and how these mechanisms adapt to both within-trial contingences, varying between pre-cue and post-cue epochs, (Gaillard et al., 2020) as well as to across task contingencies, varying as a function of task structure (Gaillard et al., 2020; Mouille et al., 2025). The present findings extend these principles to non-invasive human MEG, providing direct evidence that dynamic attentional codes can be measured at the whole-brain level from different tasks. We additionally show that both trial structure (time from target onset) and task structure (cue reliability and switching rules), systematically modulates attention decoding accuracy. The task-related differences in decoding performance provide the first line of evidence that the MEG-derived attentional code reflects behaviorally meaningful neural dynamics rather than generic signal variance. Decoding accuracy systematically decreased as cue validity was reduced, mirroring the decline in behavioral performance under uncertainty. This graded modulation indicates that the strength and spatial precision of the decoded signal scale with the reliability of top-down predictions about target location. In the 100 Valid task, participants could rely on stable expectations, resulting in highly focused pre-target attention and stronger classifier separability between left- and right-attended trials. In contrast, the 90 Valid tasks required maintaining flexible attentional sets and greater monitoring of unattended locations, producing more variable and spatially diffuse decoding traces. Under reduced cue reliability, this temporal gain was attenuated, indicating that uncertainty not only broadens the spatial distribution of attention but also dampens its temporal precision. This correspondence between decoding accuracy and behavioral efficiency aligns with reports that neural discriminability of attentional states predicts perceptual precision and decision speed (Bae and Luck, 2018; Foster et al., 2017; Wyart et al., 2015). Likewise, in all tasks, attention decoding accuracy was higher close to target presentation than earlier on in the trial, reflecting the progressive focusing of the attentional spotlight as temporal expectations build up. This temporal sharpening of decoding mirrors the anticipatory enhancement of perceptual sensitivity observed in behavioral and electrophysiological studies of attention (Nobre and van Ede, 2018; Rohenkohl and Nobre, 2011), and suggests that the decoded MEG signal captures both the spatial and temporal dynamics of top-down preparatory control. Together, these results establish that the MEG decoding signal captures task-dependent fluctuations in attentional engagement and thus constitutes a behaviorally relevant neural marker of attention.

### Single-trial tracking and direct neural–behavioral coupling

Beyond mean decoding accuracy, the Distance-to-Truth (DTT) metric and single-trial temporal trajectories provide a more sensitive, second line of evidence linking the decoded attentional signal to behavioral performance. DTT analyses revealed that attention progressively converged toward the cued target as stimulus onset approached, but that this pre-target sharpening weakened when cue reliability decreased—indicating a shift from proactive, focused orienting to a more exploratory sampling mode under uncertainty. Importantly, the magnitude of these task-related differences in attentional dynamics across participants predicted corresponding differences in behavioral performance: subjects who showed stronger differences in the spatial distribution of decoded attentional traces between tasks also exhibited greater behavioral variations in accuracy and reaction time. These neural–behavioral correlations demonstrate that trial-by-trial fluctuations in decoding strength directly translate into perceptual outcomes, supporting the notion that the decoded trajectories capture genuine attentional deployment rather than classifier noise. Such single-trial coupling between attention decoding and performance parallels intracranial observations in macaques showing that momentary deviations of the decoded attentional spotlight predict behavioral lapses (Amengual et al., 2022; Gaillard et al., 2021) as well as uncontrolled responses to task-irrelevant events (Astrand et al., 2016, 2020; Di Bello et al., 2022; Gaillard et al., 2020). All this taken together reinforces the functional validity of MEG-based attention tracking and underscores its potential for real-time applications such as neurofeedback or adaptive brain–computer interfaces.

### Rhythmic organization of attentional traces

The spectral decomposition of the decoding-derived attention trajectories provides converging evidence that spatial attention operates rhythmically in the human brain. Across all task variants, we identified robust oscillatory components within the alpha band (∼8–12 Hz), consistent with the notion that attention samples space through rhythmic cycles of enhancement and suppression (Fiebelkorn and Kastner, 2019; Gaillard et al., 2020; Landau and Fries, 2012; VanRullen, 2016). This rhythmic modulation of the decoded signal demonstrates that the neural code underlying the spatial deployment of attention is intrinsically oscillatory, even when task contingencies vary, as already reported intracortically by (Gaillard et al., 2020). The persistence of this alpha rhythmicity across tasks indicates that rhythmic sampling represents a core temporal organizing principle of attentional control, robust to changes in cue reliability and switching rules. Yet, the relative amplitude of these oscillations differed across tasks, suggesting that the strength of rhythmic modulation is flexibly tuned by top-down demands. Contrary to our initial prediction of stronger alpha power in the 90 Valid-CC condition—where continuous monitoring of the unattended side was required—alpha rhythmicity peaked in the 90 Valid-IC task, possibly reflecting stronger endogenous control to maintain and update attentional focus when unexpected cue reversals occurred in the attended hemifield. This pattern aligns with proposals that alpha oscillations index top-down inhibitory gating and flexible re-allocation of resources (Bonnefond and Jensen, 2025; Jensen, 2024; Jensen and Mazaheri, 2010; Klimesch, 2012). Importantly, these findings demonstrate that the rhythmic structure of the decoded MEG traces captures a fundamental aspect of attentional dynamics previously observed in invasive primate recordings (Gaillard et al., 2020) and human behavioral performance (Fiebelkorn et al., 2018; Landau et al., 2015).

Importantly, because alpha power distributions depended strongly on sensor selection, the observed rhythmic content may partly reflect the geometry and sensitivity profile of the occipitoparietal sensors most responsive to alpha-band fluctuations. This raises the critical question of whether the rhythmic dynamics identified here originate from localized cortical generators or from distributed large-scale coupling between parietal and frontal regions. Addressing this will require reconstructing attention-decoding trajectories at the source level, where oscillatory content can be mapped onto anatomically and functionally defined regions. Investigating rhythmic decoding from source-localized activity will be essential to establish a direct link between the temporal structure of MEG-decoded attention and the oscillatory control mechanisms that coordinate the fronto-parietal attention network (Corbetta and Shulman, 2002; Fiebelkorn and Kastner, 2019). By integrating temporal decoding and spatial source reconstruction, future work can determine how distinct cortical nodes contribute to the rhythmic sampling of space, thereby bridging non-invasive attention decoding with mechanistic models of oscillatory network control.

Overall, these findings demonstrate that MEG can capture dynamic, task-dependent fluctuations in spatial attention that parallel those observed in non-human primates. The results reveal that the structure of attentional demands reshapes the neural code for attention, modulates rhythmic sampling, and influences behavioral efficiency. By bridging invasive primate and non-invasive human research, this study provides a mechanistic link between cortical attentional control and behavior, and establishes MEG decoding of attention as a promising tool for mechanistic and clinical applications, including neurofeedback and attention-related interventions.

## Author Contributions

Conceptualization: M.B., S.B.H.

Funding: M.B., S.B.H.

Project supervision: M.B., S.B.H.

Task Design: T.C., J.M., M.B.

Preprocessing: M.F., D.S., M.M.

MEG acquisitions: T.C., M.F., S.D., D.S.

Technical development: S.D., D.S.

MEG data analysis: M.M., M.B., S.B.H.

Figure design: M.M., S.B.H.

Writing (original draft): M.M., M.B., S.B.H.

Review and editing: M.M., M.B., S.B.H.

## Data and Code Availability

Data and code will be made available open access upon publication.

## Supplementary Figures

**Figure S1.**
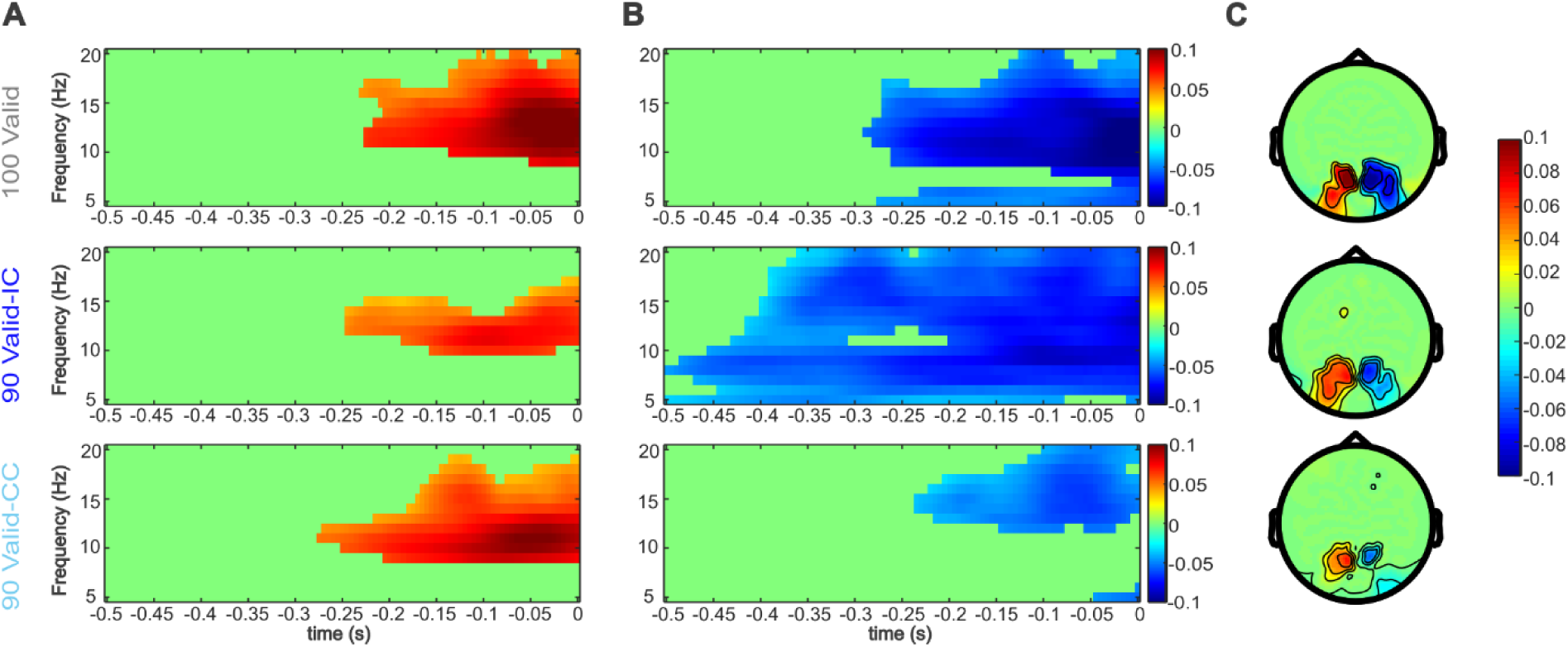
Pre-target alpha-band lateralization across tasks. Each row corresponds to one of the three tasks (100 Valid, 90 Valid-IC, 90 Valid-CC). Columns show: (A) Time–frequency representation of the Left – Right trial contrast, averaged over left-hemisphere visual channels. (B) The same Left – Right contrast, averaged over right-hemisphere visual channels. (C) Scalp topography marking the channels that exhibit significant alpha-band lateralization in the pre-target interval (–100 to 0 ms; 11–13 Hz).

**Figure S2.**
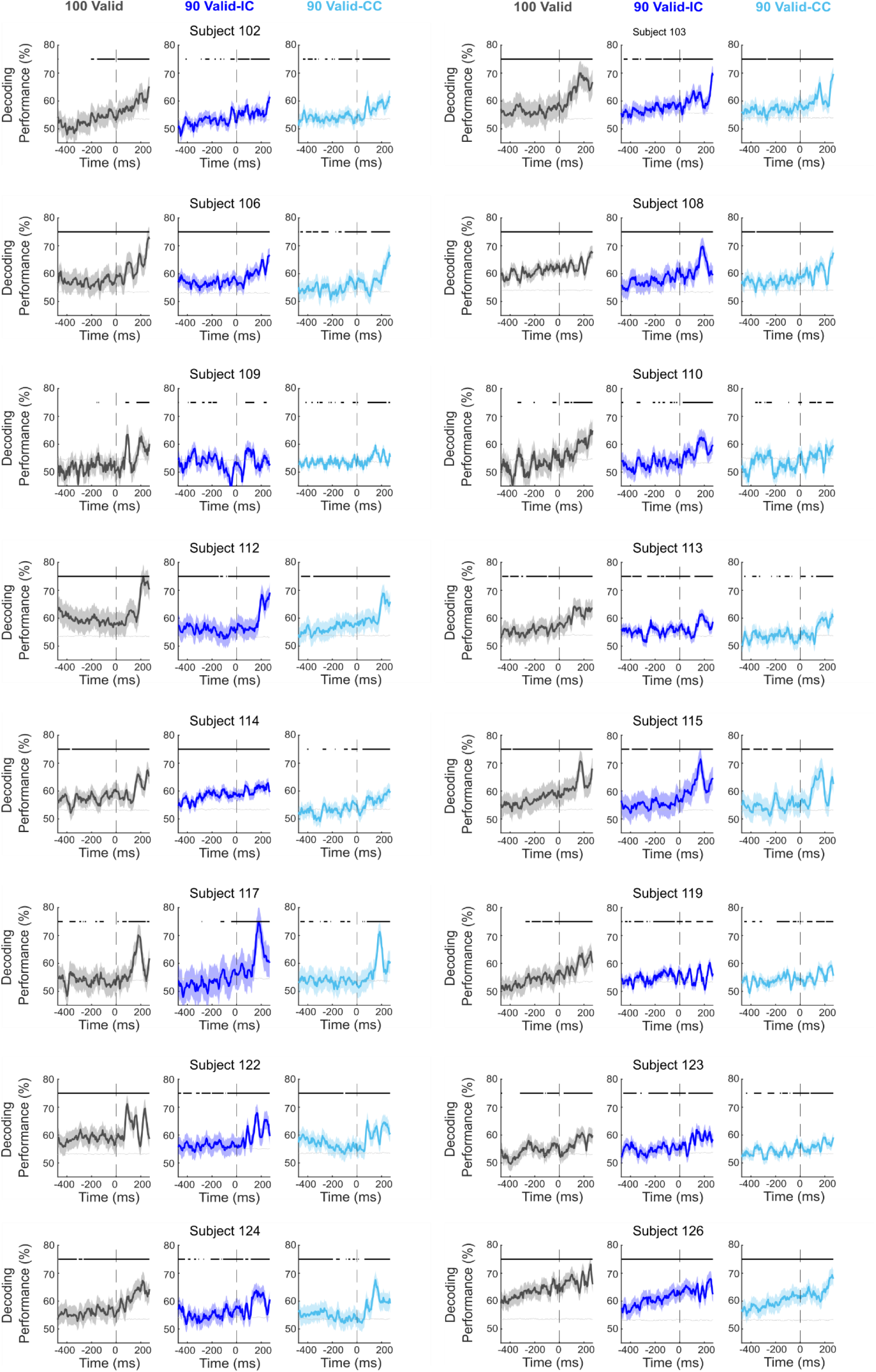
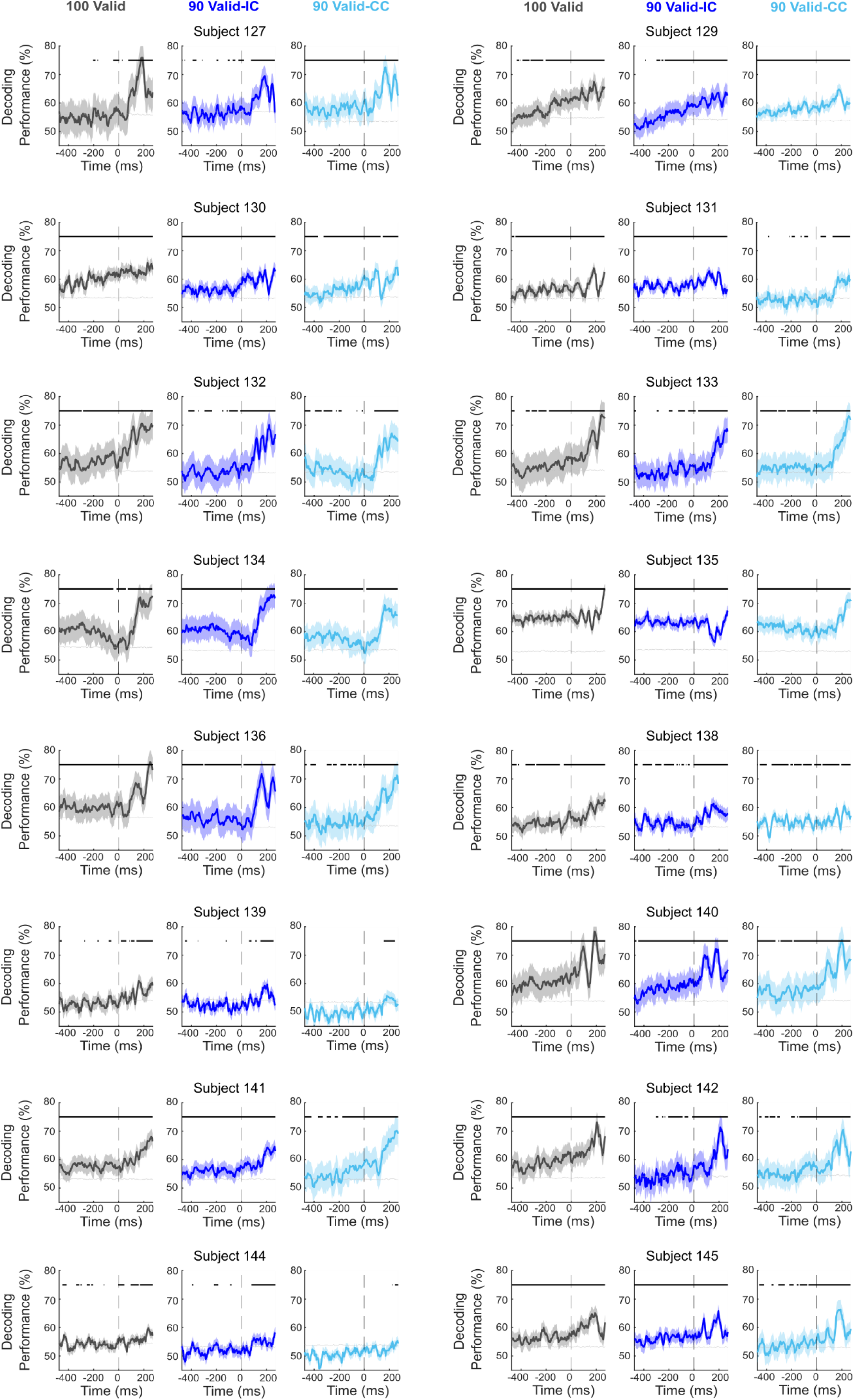
Time-resolved decoding of all subjects for visual channels. All else as in Figure 2C.

**Figure S3.**
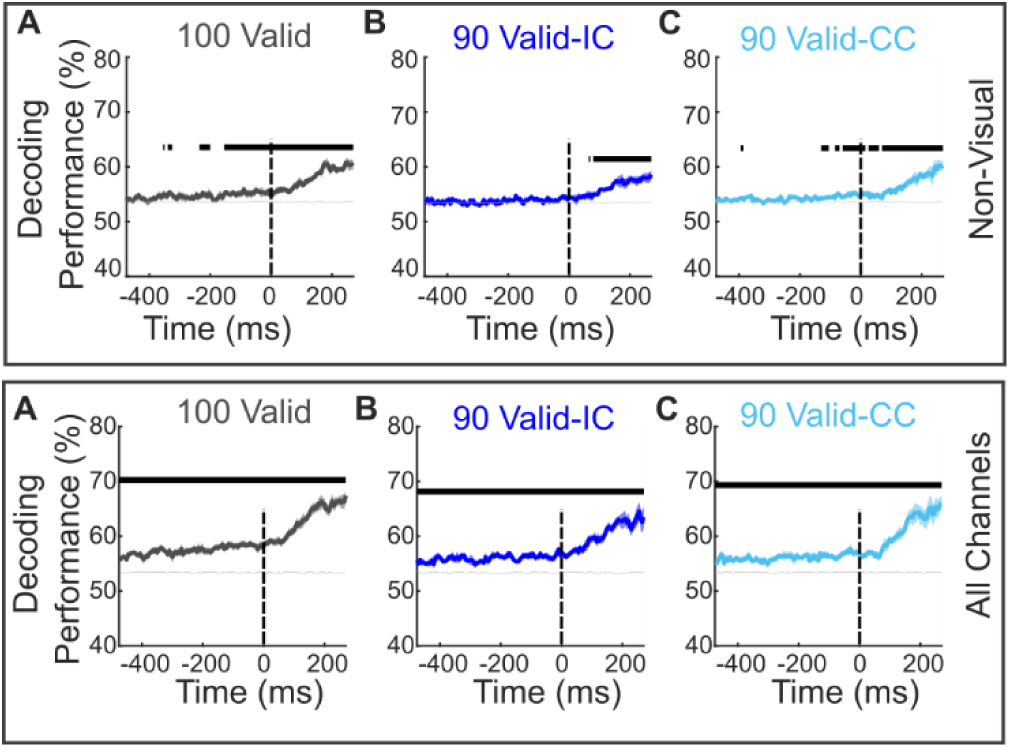
Time-resolved decoding of non-visual channels (top panel) and all channels (bottom panel) group-level results. (A) 100 Valid task (B) 90 Valid-IC task. (C) 90 Valid-CC task.

**Figure S4.**
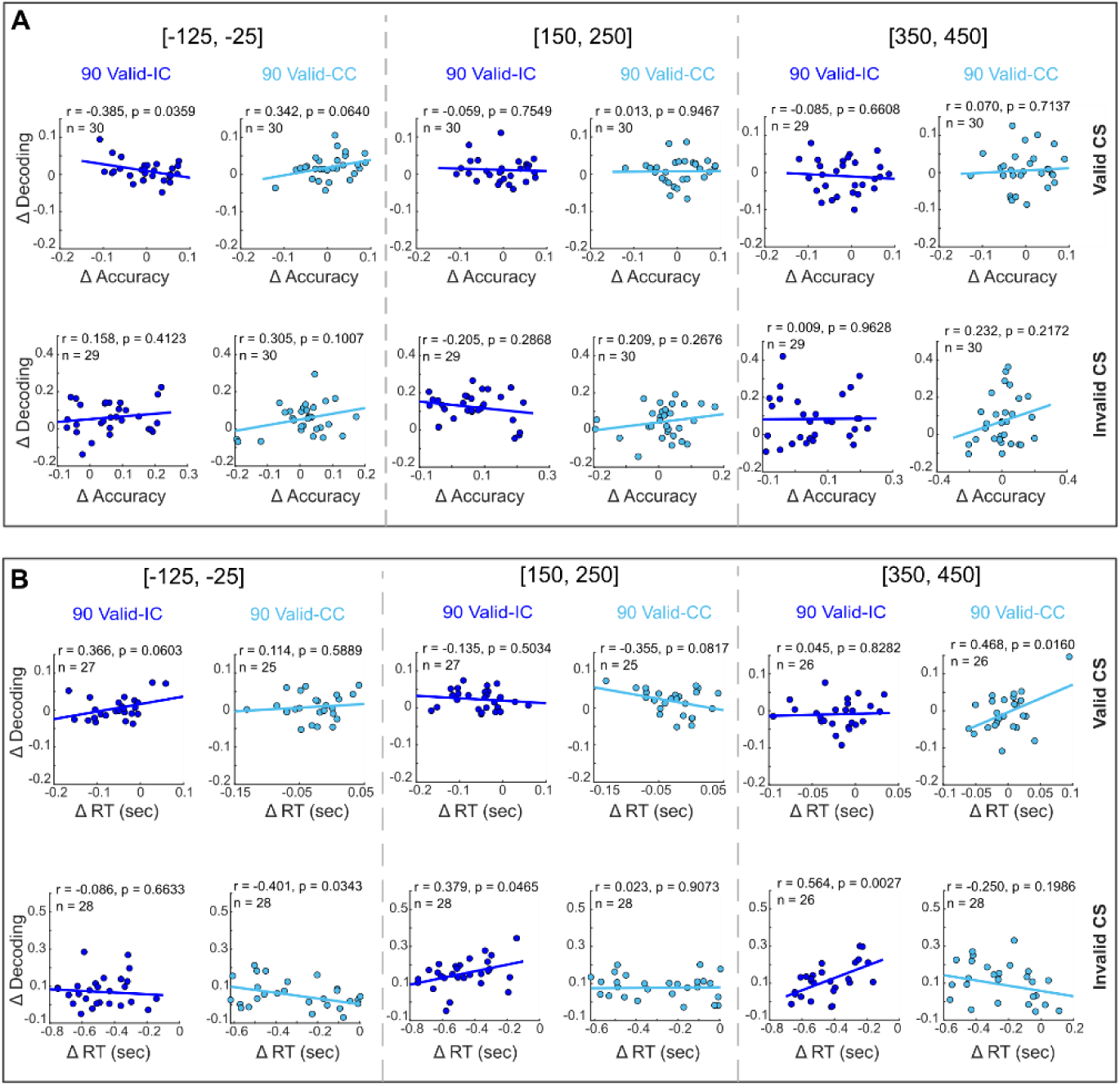
Correlation between decoding and behavioral differences across tasks. (A) **Top panel.** Correlation between changes in decoding accuracy and behavioral performance (accuracy) across participants for the Difficult–CS condition for valid trials. Bottom panel. Correlation between changes in decoding accuracy and behavioral performance (accuracy) across participants for the Difficult–CS condition for invalid trials. (B) **Top panel**. Correlation between changes in decoding accuracy and reaction RT for the **Easy–CS** condition for valid trials. **Bottom panel.** Correlation between changes in decoding accuracy and reaction RT for the **Easy–CS** condition for invalid trials. For each participant, differences were computed **relative to the 100 Valid task**, using it as a reference for valid and invalid trials from the **90 Valid-IC** and **90 Valid-CC** tasks. Correlations are shown for three temporal decoding windows ([-125, −25], [150, 250], [350, 450] ms). Each dot represents one participant; **solid lines** denote least-squares linear fits. Pearson’s *r*, *p*-values, and sample sizes (*n*) are reported within each panel. Participants with accuracy below 50% or RTs exceeding ±2 s.d. of the median in valid trials were excluded from the respective analyses.

**Figure S5.**
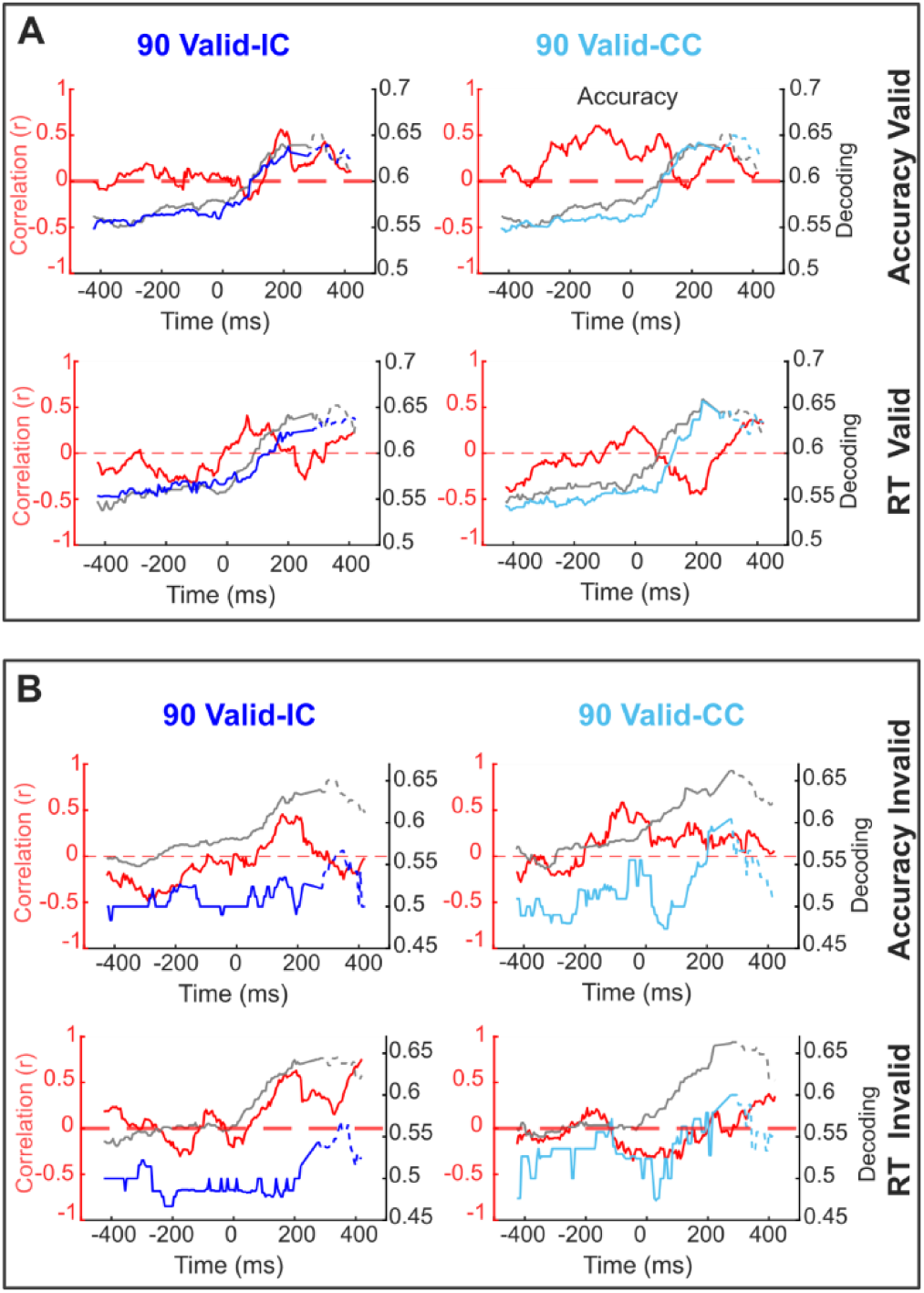
Correlation traces between decoding and behavioral differences across tasks for Valid and Invalid trials. (A) **Valid trials.** Top panels: Correlation traces in red color between changes in decoding accuracy and behavioral performance (accuracy) across participants for the Difficult–IS condition for 90 Valid-IC and 90 Valid-CC tasks with showing decoding trace for 100 Valid in gray color as reference and 90 Valid-IC decoding trace in dark blue (left_panel) and light blue for 90 Valid-CC (right panel). Bottom panel: Same as top panel, but the correlation traces are between changes in decoding accuracy and RT across participants for the Easy–IS condition. (B) **Invalid trials.** Top panel and bottom panel are the same as (A), but for invalid trials. For each participant, differences were computed relative to the 100 Valid task, using it as a reference for valid and invalid trials in the 90 Valid-IC and 90 Valid-CC tasks. Each time point in the correlation trace represents the correlation of 100 ms sliding-window decoding accuracy values with behavioral differences across participants.

**Figure S6.**
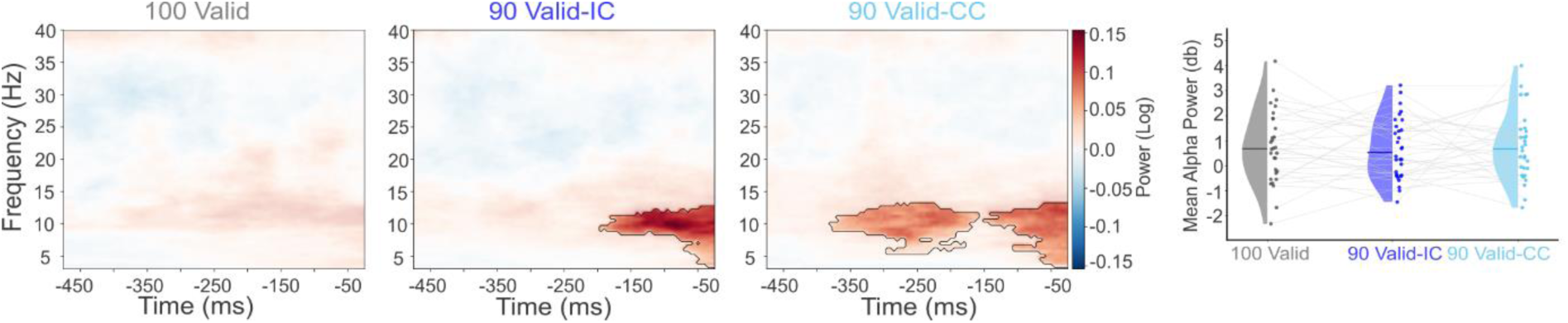
**Time-frequency representation (TFR) of DTT time series non-visual channels. (A) Left panel**, Contrast TFR for the 100 Valid task, showing significant clusters (with non-significant areas displayed with reduced transparency). **Middle panel**, Contrast TFR for the 90 Valid-IC task, with significant clusters. **Right Panel**, Contrast TFR for the 90 Valid-CC task, with significant clusters.

**Figure S7.**
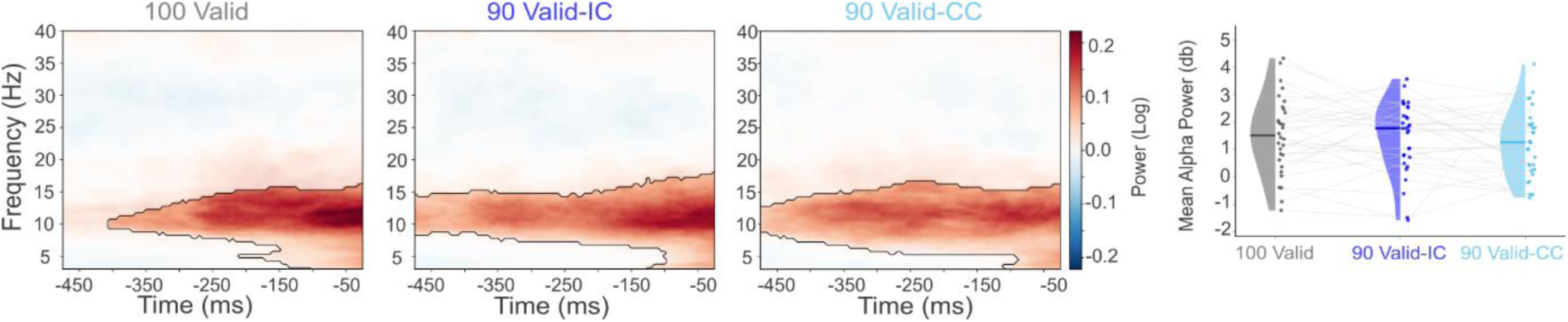
**Time-frequency representation (TFR) of DTT time series all channels. (A) Left panel**, Contrast TFR for the 100 Valid task, showing significant clusters (with non-significant areas displayed with reduced transparency). **Middle panel**, Contrast TFR for the 90 Valid-IC task, with significant clusters. **Right Panel**, Contrast TFR for the 90 Valid-CC task, with significant clusters.

## References

Amengual, J.L., Di Bello, F., Ben Hadj Hassen, S., Ben Hamed, S., 2022a. Distractibility and impulsivity neural states are distinct from selective attention and modulate the implementation of spatial attention. Nat Commun 13, 4796. 10.1038/s41467-022-32385-y

Amengual, J.L., Di Bello, F., Ben Hadj Hassen, S., Ben Hamed, S., 2022b. Distractibility and impulsivity neural states are distinct from selective attention and modulate the implementation of spatial attention. Nat Commun 13, 4796. 10.1038/s41467-022-32385-y

Astrand, E., Wardak, C., Baraduc, P., Ben Hamed, S., 2016. Direct Two-Dimensional Access to the Spatial Location of Covert Attention in Macaque Prefrontal Cortex. Current Biology 26, 1699–1704. 10.1016/j.cub.2016.04.054

Astrand, E., Wardak, C., Ben Hamed, S., 2020. Neuronal population correlates of target selection and distractor filtering. NeuroImage 209, 116517. 10.1016/j.neuroimage.2020.116517

Bae, G.-Y., Luck, S.J., 2018. Dissociable Decoding of Spatial Attention and Working Memory from EEG Oscillations and Sustained Potentials. J. Neurosci. 38, 409–422. 10.1523/JNEUROSCI.2860-17.2017

Bonnefond, M., Jensen, O., 2025. The role of alpha oscillations in resisting distraction. Trends in Cognitive Sciences 29, 368–379. 10.1016/j.tics.2024.11.004

Cichy, R.M., Pantazis, D., Oliva, A., 2014. Resolving human object recognition in space and time. Nat Neurosci 17, 455–462. 10.1038/nn.3635

Corbetta, M., Shulman, G.L., 2002a. Control of goal-directed and stimulus-driven attention in the brain. Nat Rev Neurosci 3, 201–215. 10.1038/nrn755

Corbetta, M., Shulman, G.L., 2002b. Control of goal-directed and stimulus-driven attention in the brain. Nat Rev Neurosci 3, 201–215. 10.1038/nrn755

Crouzet, S.M., VanRullen, R., 2017. The rhythm of attentional stimulus selection during visual competition (preprint). Neuroscience. 10.1101/105239

De Sousa, C., Gaillard, C., Di Bello, F., Ben Hadj Hassen, S., Ben Hamed, S., 2021. Behavioral validation of novel high resolution attention decoding method from multi-units & local field potentials. NeuroImage 231, 117853. 10.1016/j.neuroimage.2021.117853

Di Bello, F., Ben Hadj Hassen, S., Astrand, E., Ben Hamed, S., 2022a. Prefrontal Control of Proactive and Reactive Mechanisms of Visual Suppression. Cerebral Cortex 32, 2745–2761. 10.1093/cercor/bhab378

Di Bello, F., Ben Hadj Hassen, S., Astrand, E., Ben Hamed, S., 2022b. Prefrontal Control of Proactive and Reactive Mechanisms of Visual Suppression. Cerebral Cortex 32, 2745–2761. 10.1093/cercor/bhab378

Donoghue, T., Haller, M., Peterson, E.J., Varma, P., Sebastian, P., Gao, R., Noto, T., Lara, A.H., Wallis, J.D., Knight, R.T., Shestyuk, A., Voytek, B., 2020. Parameterizing neural power spectra into periodic and aperiodic components. Nature Neuroscience 23, 1655–1665. 10.1038/s41593-020-00744-x

Fiebelkorn, I.C., Kastner, S., 2019a. A Rhythmic Theory of Attention. Trends in Cognitive Sciences 23, 87–101. 10.1016/j.tics.2018.11.009

Fiebelkorn, I.C., Kastner, S., 2019b. A Rhythmic Theory of Attention. Trends in Cognitive Sciences 23, 87–101. 10.1016/j.tics.2018.11.009

Fiebelkorn, I.C., Pinsk, M.A., Kastner, S., 2018. A Dynamic Interplay within the Frontoparietal Network Underlies Rhythmic Spatial Attention. Neuron 99, 842–853.e8. 10.1016/j.neuron.2018.07.038

Fiebelkorn, I.C., Saalmann, Y.B., Kastner, S., 2013. Rhythmic Sampling within and between Objects despite Sustained Attention at a Cued Location. Current Biology 23, 2553–2558. 10.1016/j.cub.2013.10.063

Foster, J.J., Sutterer, D.W., Serences, J.T., Vogel, E.K., Awh, E., 2017. Alpha-Band Oscillations Enable Spatially and Temporally Resolved Tracking of Covert Spatial Attention. Psychol Sci 28, 929–941. 10.1177/0956797617699167

Gaillard, C., Ben Hadj Hassen, S., Di Bello, F., Bihan-Poudec, Y., VanRullen, R., Ben Hamed, S., 2020a. Prefrontal attentional saccades explore space rhythmically. Nat Commun 11, 925. 10.1038/s41467-020-14649-7

Gaillard, C., Ben Hadj Hassen, S., Di Bello, F., Bihan-Poudec, Y., VanRullen, R., Ben Hamed, S., 2020b. Prefrontal attentional saccades explore space rhythmically. Nat Commun 11, 925. 10.1038/s41467-020-14649-7

Gaillard, C., Ben Hamed, S., 2022. The neural bases of spatial attention and perceptual rhythms. Eur J of Neuroscience 55, 3209–3223. 10.1111/ejn.15044

Gaillard, C., Sousa, C.D., Amengual, J., Loriette, C., Ziane, C., Hassen, S.B.H., Bello, F.D., Hamed, S.B., 2021. Attentional brain rhythms during prolonged cognitive activity. 10.1101/2021.05.26.445730

Grootswagers, T., Wardle, S.G., Carlson, T.A., 2017. Decoding Dynamic Brain Patterns from Evoked Responses: A Tutorial on Multivariate Pattern Analysis Applied to Time Series Neuroimaging Data. J Cogn Neurosci 29, 677–697. 10.1162/jocn_a_01068

Hamed, S.B., 2025. Decoding Covert Visual Attention in Space and Time from Neural Signals. Annual Review of Vision Science 11, 495–520. 10.1146/annurev-vision-101322-011902

Holcombe, A.O., Chen, W.-Y., 2013. Splitting attention reduces temporal resolution from 7 Hz for tracking one object to <3 Hz when tracking three. Journal of Vision 13, 12–12. 10.1167/13.1.12

Jaiswal, A., Nenonen, J., Stenroos, M., Gramfort, A., Dalal, S.S., Westner, B.U., Litvak, V., Mosher, J.C., Schoffelen, J.-M., Witton, C., Oostenveld, R., Parkkonen, L., 2020. Comparison of beamformer implementations for MEG source localization. NeuroImage 216, 116797. 10.1016/j.neuroimage.2020.116797

Jensen, O., 2024. Distractor inhibition by alpha oscillations is controlled by an indirect mechanism governed by goal-relevant information. Commun Psychol 2, 36. 10.1038/s44271-024-00081-w

Jensen, O., Mazaheri, A., 2010. Shaping Functional Architecture by Oscillatory Alpha Activity: Gating by Inhibition. Front. Hum. Neurosci. 4. 10.3389/fnhum.2010.00186

Kienitz, R., Schmiedt, J.T., Shapcott, K.A., Kouroupaki, K., Saunders, R.C., Schmid, M.C., 2018. Theta Rhythmic Neuronal Activity and Reaction Times Arising from Cortical Receptive Field Interactions during Distributed Attention. Current Biology 28, 2377–2387.e5. 10.1016/j.cub.2018.05.086

King, J.-R., Dehaene, S., 2014. Characterizing the dynamics of mental representations: the temporal generalization method. Trends in Cognitive Sciences 18, 203–210. 10.1016/j.tics.2014.01.002

Klimesch, W., 2012. Alpha-band oscillations, attention, and controlled access to stored information. Trends in Cognitive Sciences 16, 606–617. 10.1016/j.tics.2012.10.007

Konttila, T., Mäntynen, V., Stenroos, M., 2014. Comparison of minimum-norm estimation and beamforming in electrocardiography with acute ischemia. Physiol. Meas. 35, 623. 10.1088/0967-3334/35/4/623

Landau, A.N., Fries, P., 2012. Attention Samples Stimuli Rhythmically. Current Biology 22, 1000–1004. 10.1016/j.cub.2012.03.054

Landau, A.N., Schreyer, H.M., van Pelt, S., Fries, P., 2015. Distributed Attention Is Implemented through Theta-Rhythmic Gamma Modulation. Current Biology 25, 2332–2337. 10.1016/j.cub.2015.07.048

Maris, E., Oostenveld, R., 2007. Nonparametric statistical testing of EEG- and MEG-data. Journal of Neuroscience Methods 164, 177–190. 10.1016/j.jneumeth.2007.03.024

Michel, R., Dugué, L., Busch, N.A., 2020. Perceptual rhythms are driven by oscillations in visual precision. Journal of Vision 20, 1164. 10.1167/jov.20.11.1164

Moca, V.V., Bârzan, H., Nagy-Dăbâcan, A., Mureșan, R.C., 2021. Time-frequency super-resolution with superlets. Nat Commun 12, 337. 10.1038/s41467-020-20539-9

Mouille, A., Gaillard, C., Astrand, E., Wardak, C., Amengual, J.L., Hamed, S.B., 2025. The prefrontal cortex encodes task-identity information and flexibly adjusts its sensory processes as a function of the specific ongoing task. PLOS Biology 23, e3003353. 10.1371/journal.pbio.3003353

Nobre, A.C., van Ede, F., 2018. Anticipated moments: temporal structure in attention. Nat Rev Neurosci 19, 34–48. 10.1038/nrn.2017.141

Rohenkohl, G., Nobre, A.C., 2011. Alpha Oscillations Related to Anticipatory Attention Follow Temporal Expectations. J. Neurosci. 31, 14076–14084. 10.1523/JNEUROSCI.3387-11.2011

Samaha, J., Sprague, T.C., Postle, B.R., 2016. Decoding and Reconstructing the Focus of Spatial Attention from the Topography of Alpha-band Oscillations. Journal of Cognitive Neuroscience 28, 1090–1097. 10.1162/jocn_a_00955

Spyropoulos, G., Bosman, C.A., Fries, P., 2018. A theta rhythm in macaque visual cortex and its attentional modulation. Proc. Natl. Acad. Sci. U.S.A. 115. 10.1073/pnas.1719433115

Thaler, L., Schütz, A.C., Goodale, M.A., Gegenfurtner, K.R., 2013. What is the best fixation target? The effect of target shape on stability of fixational eye movements. Vision Research 76, 31–42. 10.1016/j.visres.2012.10.012

Thut, G., Nietzel, A., Brandt, S.A., Pascual-Leone, A., 2006. α-Band Electroencephalographic Activity over Occipital Cortex Indexes Visuospatial Attention Bias and Predicts Visual Target Detection. J. Neurosci. 26, 9494–9502. 10.1523/JNEUROSCI.0875-06.2006

Ungerleider, S.K.A.L.G., 2000. Mechanisms of Visual Attention in the Human Cortex. Annu. Rev. Neurosci. 23, 315–341. 10.1146/annurev.neuro.23.1.315

VanRullen, R., 2018. Attention Cycles. Neuron 99, 632–634. 10.1016/j.neuron.2018.08.006

VanRullen, R., 2016. Perceptual Cycles. Trends in Cognitive Sciences 20, 723–735. 10.1016/j.tics.2016.07.006

VanRullen, R., 2013. Visual Attention: A Rhythmic Process? Current Biology 23, R1110–R1112. 10.1016/j.cub.2013.11.006

Worden, M.S., Foxe, J.J., Wang, N., Simpson, G.V., 2000. Anticipatory Biasing of Visuospatial Attention Indexed by Retinotopically Specific α-Bank Electroencephalography Increases over Occipital Cortex. J. Neurosci. 20, RC63–RC63. 10.1523/JNEUROSCI.20-06-j0002.2000

Wyart, V., Myers, N.E., Summerfield, C., 2015. Neural Mechanisms of Human Perceptual Choice Under Focused and Divided Attention. J. Neurosci. 35, 3485–3498. 10.1523/JNEUROSCI.3276-14.2015

